# Exploring amino acid functions in a deep mutational landscape

**DOI:** 10.1101/2020.05.26.116756

**Authors:** Alistair Dunham, Pedro Beltrao

## Abstract

Amino acids fulfil a diverse range of roles in proteins, each utilising its chemical properties in different ways in different contexts to create required functions. For example, cysteines form disulphide or hydrogen bonds in different circumstances and charged amino acids do not always make use of their charge. The repertoire of amino acid functions and the frequency at which they occur in proteins remains understudied. Measuring large numbers of mutational consequences, which can elucidate the role an amino acid plays, was prohibitively time consuming until recent developments in deep mutational scanning. In this study we gathered data from 28 deep mutational scanning studies, covering 6291 positions in 30 proteins, and used the consequences of mutation at each position to define a mutational landscape. We demonstrated rich relationships between this landscape and biophysical or evolutionary properties. Finally, we identified 100 functional amino acid subtypes with a data-driven clustering analysis and studied their features, including their frequencies and chemical properties such as tolerating polarity, hydrophobicity or being intolerant of charge or specific amino acids. The mutational landscape and amino acid subtypes provide a foundational catalogue of amino acid functional diversity, which will be refined as the number of studied protein positions increases.

## Introduction

Amino acids fulfil a diverse range of roles in proteins, each utilising its chemical properties in different ways in different contexts in order to create required protein functions. Classically, amino acids split into five broad groups based on their physicochemistry; aliphatic, aromatic, polar, positively charged and negatively charged. In addition proline and glycine uniquely form a backbone loop and lack a side chain, respectively. However, the same amino acid can have different effects in different contexts. For example, consider three tyrosine contexts; Tyr197 in the active site of NADPH dehydrogenase, a tyrosine phosphosite and Tyr198 in inositol polyphosphate multikinase. The first two use the hydroxyl group in divergent ways; forming a stabilising hydrogen bond to the enzymes substrate and covalently bonding to the new phosphate group during phosphorylation. The final example uses a completely different property, the aromatic ring, to form a stabilising interaction with a sulphur atom in a nearby methionine (Weber and Warren, 2019). Even in the narrow context of catalytic sites the same amino acids have been shown to perform a range of active roles across the proteome (Ribeiro et al., 2020). The role an amino acid fills at a given position is defined by its chemical environment, which is determined by many factors including nearby amino acids, secondary structure, post-translational modifications, and bound ligands. Taking account of this environment helps predict whether a site is functional, thus demonstrating the environment’s importance (Rice and Eisenberg, 1997; Torng and Altman, 2019).

An amino acid’s role affects the consequences of different substitutions at that position, known as the positions mutational profile. Evolutionary conservation data confirms this, with models of substitution likelihood that account for protein context performing better than universal models (Huang and Bystroff, 2006; Müller et al., 2001; Rice and Eisenberg, 1997). Similarly, on a gene by gene level, alanine scanning experiments have shown the power of mutations to infer positional functions in proteins using mutations, for example mapping hGH receptor interactions (Cunningham and Wells, 1989) and the CD4 binding site (Ashkenazi et al., 1990). This association between mutational consequences and an amino acid’s contextual properties is biologically important, both for understanding protein chemistry and in predicting the consequences of mutations.

Given the strong link between a position’s structural and functional role and mutational consequences, the landscape of mutational profiles across the proteome can be used for an unbiased, proteome wide exploration of the functional diversity of amino acid roles. Until recently experimentally characterising this landscape was difficult, but the new deep mutational scanning technique (Fowler and Fields, 2014) measures mutational profiles at very high throughput. These experiments directly measure fitness for all possible variants in a protein or region by generating a comprehensive mutant library and subjecting it to selection where survival is dependent on the target protein’s function. Sequencing the library before and after selection determines the fitness of variants by measuring whether they have been enriched or depleted relative to the wild type during selection. Data from these experiments has been used to explore properties of the genes being studied, to determine general properties of insertion types (Gray et al., 2017) and for training variant effect predictors (Gray et al., 2018).

Here we explore the combined mutational landscape of 30 genes using 33 deep mutational scans from 28 studies. We show that multiple datasets can be meaningfully combined and use them to explore the diverse functions of each amino acid. First, suitable studies were selected and their scores normalised to a common scale. Second, we derived and analysed their combined mutational landscape, relating it to a range of biophysical properties. Finally we explored the diversity of amino acid roles using the mutational landscape, clustering each amino acid into typical subtypes and demonstrating these subtypes represent meaningful biological groups.

## Results

### A unified deep mutational scanning landscape

The first step to utilising a dataset drawn from many deep mutational scanning studies is combining them in a meaningful way, allowing scores to be compared. We selected 33 deep mutational scans from 28 studies covering 6291 positions in 30 proteins across nine species (**figure 1A, table S1**). These proteins have a broad range of structures and functions. We required that studies applied selection pressures directly related to natural function and that their fitness scores could be transformed to a common scale. To prepare each study (**figure 1B**) we first averaged replicates and either selected the most appropriate condition or averaged comparable ones. Some studies generated sequences with multiple variants, so we averaged across variants containing each substitution, when individual substitutions were not measured explicitly (**figure S1**). Next, substitution fitness scores were transformed into a common enrichment ratio (ER) score, which measures the enrichment of a variant during selection relative to the change in the wild type. Thus, positive ER scores mean the variant is advantageous, zero is neutral and negative scores deleterious. We normalised scores from each study against the median of the lowest 10% of scores, reasoning that the worst substitutions (e.g. nonsense mutations) result in complete loss of function and are comparable between studies. We filtered positions with scores for less than 15 of the possible 20 nonsynonymous substitutions (including nonsense) to focus on positions with sufficient data and imputed remaining missing data (see Methods). This resulted in complete sets of normalised ER scores for substitutions to all 20 amino acids at 6291 unique positions across 30 genes, with 98.16% of nonsynonymous ER scores experimentally measured. We refer to the vector of a position’s ER scores as its mutational profile and the combined mutational profiles for all positions as the mutational landscape.

**Figure 1.**
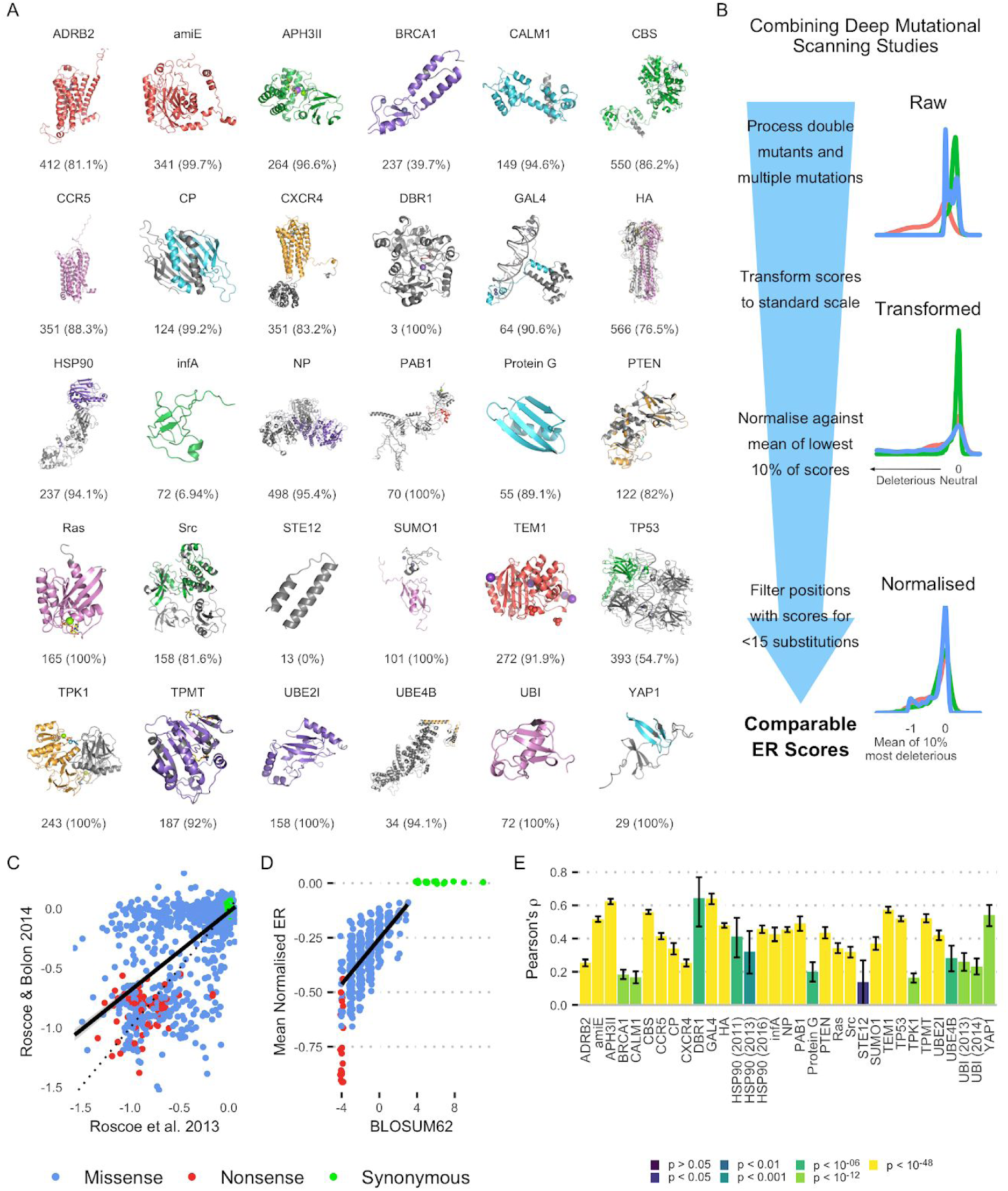
Combining deep mutational scanning studies. **A:** Proteins in the combined dataset, with the structure used, number of positions with mutational profiles and the percentage of these in the structure model. **B:** Normalisation pipeline. **C:** Correlation between ER scores from two deep mutational scans on Ubiquitin, both from the Bolon lab (*r*^2^ = 0.4676, *p* < 2.2 × 10^−16^). **D:** Relationship between mean normalised ER score for an amino acid substitution and corresponding BLOSUM62 score. The scores for missense variants strongly correlate (*r*^2^ = 0.4194, *p* < 2.2 × 10^−16^) **E:** Correlation between normalised ER score and log_10_SIFT score in each study

Three genes (HSP90, TEM1 & UBI) were covered by multiple studies (**figures 1C** & **S2**). The scores from these results were correlated (HSP90 *r*^2^ = 0.4038, TEM1 *r*^2^ = 0.994, UBI *r*^2^ = 0.4676), despite differences in the experimental conditions in HSP90 and UBI, suggesting the scores are robust and represent variants’ biological properties rather than the experimental method or selection criteria. The two TEM1 studies were so similar to each other that only the most recent was used. On the other hand the HSP90 and UBI studies were sufficiently different that they could all be retained, and used to check consistency of later results.

Deep mutational scanning results significantly relate to evolutionary conservation, which is to be expected because natural selection also samples variants and enriches them based on fitness. We validated our approach by showing this relationship was maintained in bulk statistics and for individual variants in our data. Substitution matrices, such as BLOSUM62 (Henikoff and Henikoff, 1992), measure how frequently each amino acid is mutated to any other, based on large sequence alignments. The mean ER score for each missense substitution type (A → C, A → D, …) in our dataset correlates well with BLOSUM62 (**figure 1D**, *r*^2^ = 0.4194, *p* < 2.2 × 10^−16^). BLOSUM62 also provides scores for synonymous substitutions, measuring the amino acids’ average conservation, while synonymous substitutions have 0 ER in deep mutational scans. In the mutational landscape data overall conservation is measured by the mean ER score for all substitutions away from that amino acid, which correlates with synonymous BLOSUM62 scores (*r*^2^ = 0.3191, *p* = 0.009452). Secondly, ER scores for specific substitutions correlate with log_10_SIFT scores (**figure 1E**), which summarise the likelihood of a specific substitution based on conservation (Vaser et al., 2015). The varying correlation with SIFT between studies suggests that the different experimental selection pressures vary in the degree they mirror natural evolutionary pressures.

The combined dataset gives us a large number of mutational profiles, together constituting a subset of the overall proteome mutational landscape. While it covers a relatively small subset of the vast proteome space the dataset does cover a reasonably varied and representative sample, with proteins covering a range of species, functions and environments. Thus we generated an approximation for the mutational landscape of proteins and proceeded to analyse amino acid diversity with it.

### Biophysical relationships in the deep mutational landscape

The link between mutational consequences and amino acid’s roles means the mutational landscape we have derived is a quantitative representation of the diverse roles amino acids can play in proteins. Thus, biophysical properties would be expected to map onto this space in meaningful patterns, depending on how they relate to different roles. These relationships can be visualised by dimensionality reduction of the mutational landscape using UMAP (McInnes et al., 2018) and PCA (**figure 2**), in which positions’ mutational profiles are mapped to a new space designed to have meaningful axes while maintaining the spatial proximity of positions with similar consequences. If different studies’ experimental setups strongly influenced positions’ ER profiles positions would separate by protein or study of origin. Instead, all studies span UMAP space so the dimensionality reduction demonstrates a lack of batch effects (**figure S3**). In addition, the same protein positions assayed in different studies are significantly closer together in UMAP space, on average, than random pairs of positions (**figure S4**, mean 2.105 units closer, one tailed Mann-Whitney U-Test: *p* = 1.141 × 10^−13^), which suggests the method is replicable and validates our approach to building a combined dataset.

**Figure 2.**
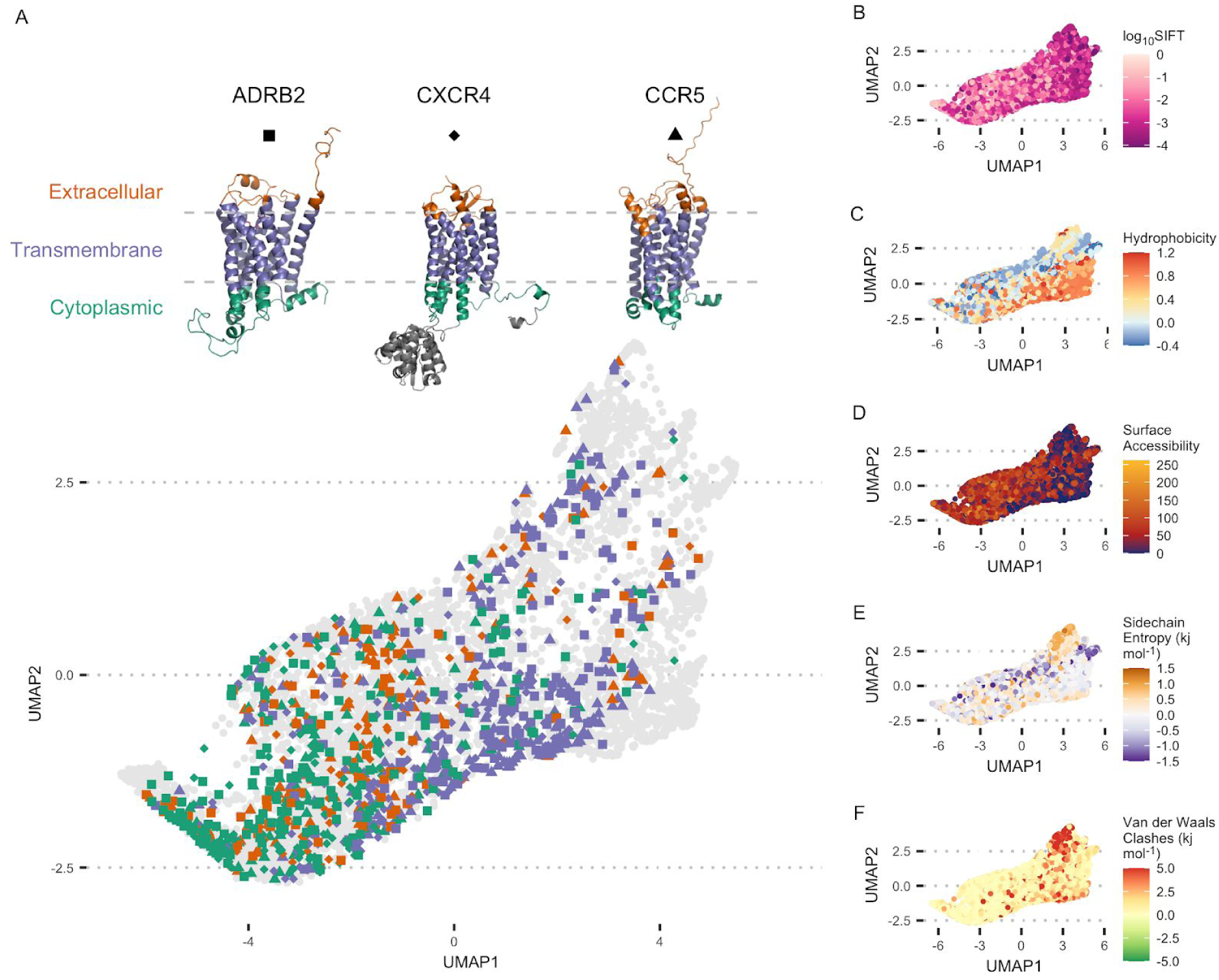
UMAP projection of the combined deep mutational landscape, coloured by different biophysical factors at each protein position. **A:** Separation of sites in the extracellular, transmembrane and cytoplasmic domains of three transmembrane proteins (ADRB2, CXCR4 and CCR5). **B:** Mean log10 SIFT score, showing conservation relates most to the first UMAP component (*r*^2^ = 0.2169, *p* < × 10^−16^). **C:** Mean amino acid hydrophobicity. **D:** All atom absolute residue surface accessibility. **E:** Mean sidechain entropy term from all FoldX substitution ΔΔ*G* predictions at a position. **F:** Mean Van der Waals clashes term from all FoldX substitution ΔΔ*G* predictions at a position.

The relationship between the mutational landscape and the biophysical properties of protein positions is most clearly illustrated by highlighting protein domains in UMAP space (**figure 2A**), and observing separation by function. In this case we highlight positions in three transmembrane proteins (ADRB2, CCR5 & CXCR4) and show they divide primarily by the solvent environment they are exposed to; either the lipid membrane or hydrophilic intra/extracellular solvent. Quantitative properties can be also demonstrated in the landscape, the strongest of which is the strong role of mean normalised ER and its link to overall evolutionary conservation at a position (**figure 2B**). The first UMAP dimension strongly correlates with mean normalised ER (*r*^2^ = 0.9036, *p* < 2.2 × 10^−16^) and more weakly with mean log_10_SIFT score (*r*^2^ = 0.2169, *p* < 2.2 × 10^−16^), which is a measure for evolutionary conservation. Indeed the first principal component (46.7% of the variance, **figure S5**) essentially is mean normalised ER (*r*^2^ = 0.9991, *p* < 2.2 × 10^−16^).

We computationally derived each position’s physical properties from its sequence and structure (**table S2**), and demonstrated strong patterns in UMAP space. Positions segregate on the mean hydrophobicity of their wild-type amino acid (Bandyopadhyay and Mehler, 2008) (**figure 2C**), and show a strongly related pattern in their surface accessibility (Hubbard and Thornton, 1993) (**figure 2D**). FoldX (Schymkowitz et al., 2005) uses a physics derived force field to estimate ΔΔ*G* changes in folding energy after mutation, breaking the result down into components from different physical sources (Van der Waals, electrostatics, etc). The average ΔΔ*G* from each component across all substitutions at a position is indicative of the physical effects of the wild-type amino acid, but with the opposite ΔΔ*G* sign because substitutions disrupt wild-type interactions. For example, positive hydrogen bond ΔΔ*G* values for most substitutions suggest that the wild-type makes stabilising hydrogen bonds that are disrupted by mutations. These measurements of positions’ physical properties also display strong patterns in UMAP space (**figure 2E** & **F**). Thus, we show that the mutational landscape has rich and complex relationships with a range of biophysical properties, and expect there to be others not highlighted in our analysis. Therefore, a position’s location in this landscape indicates the likely properties it has, creating a quantitative map of the diverse functions of amino acids in proteins. Consequently positions with similar roles or in similar environments group together in the landscape, as seen in domains of ADRB2, CCR5 and CXCR4, and so reduced dimensionality representations of the mutational landscape are a good space to evaluate the diversity of amino acid roles.

### Mapping the diversity of amino acid subtypes

Next, we used the mutational landscape to study the diverse roles played by each amino acid. We have shown that positions with similar properties, and therefore likely similar roles, cluster in this space so an approach to mapping role diversity would be to split positions of each amino acid into typical subtypes. This would give an overview of typical functions played by each amino acid, their relationships to other amino acids and how frequent they are. We clustered the mutational profiles of each amino acid’s wild-type positions independently (**figure 3A, Methods**), in order to study each amino acid’s roles in an unbiased manner.

**Figure 3.**
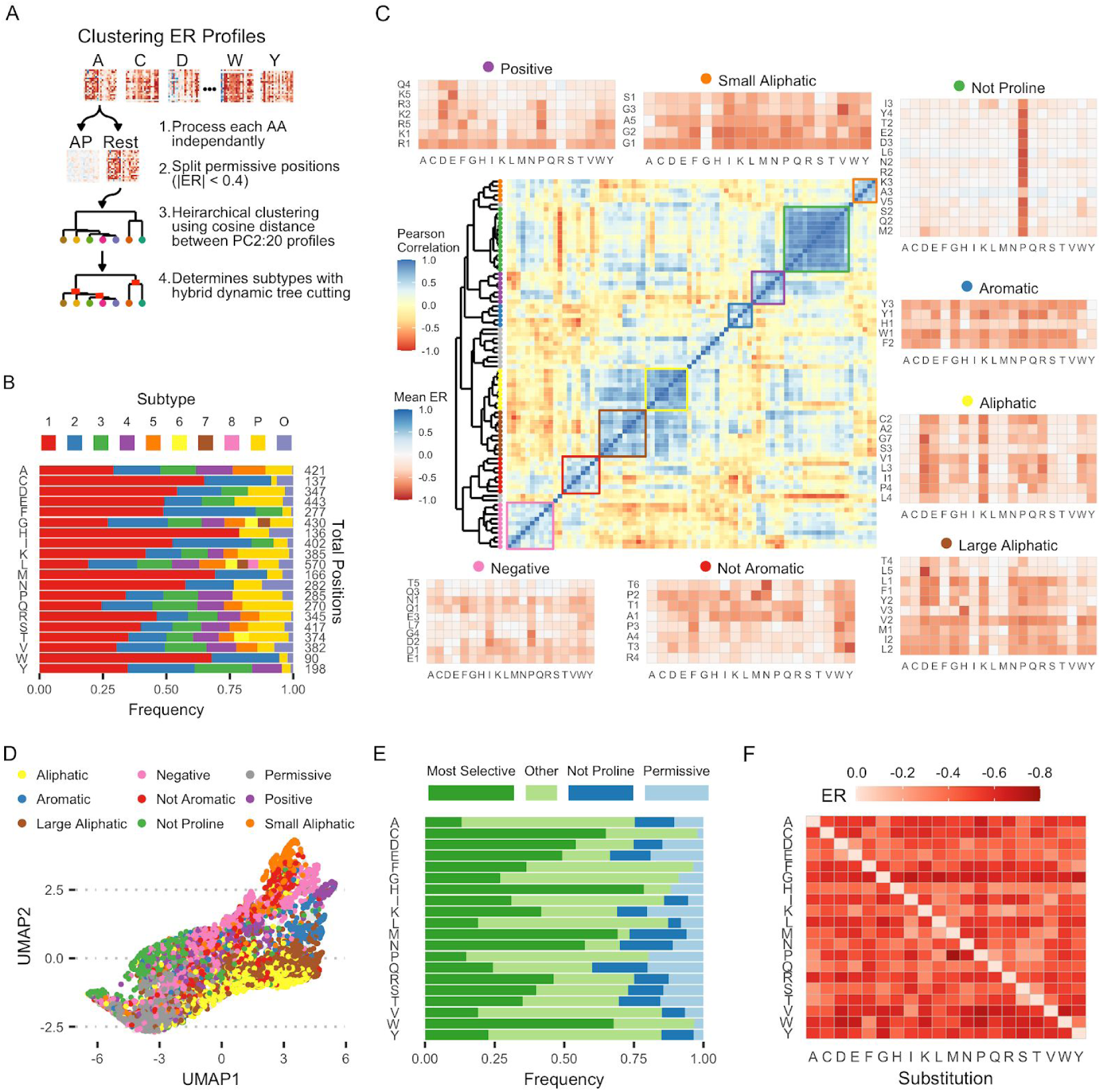
Amino acid subtypes. **A:** Subtype clustering pipeline. **B:** Frequency of each subtype and total number of positions of each amino acid. **C:** Heatmap showing the correlation between subtypes mean mutation profiles. Clusters of subtypes are colour coded with the mutational profiles of their constituent subtypes shown. **D:** Positions from each correlation cluster mapped to the UMAP projection of the deep mutational landscape. **E:** Frequency of the most selective (highest mean normalised ER), not proline (see **C**) and permissive subtypes of each amino acid. **F:** Mutational profile of the most selective subtype (see **E**) of each amino acid.

Clustering mutational profiles was dominated by the position’s mean ER score, which prevented separation by other properties. To avoid this, we clustered using the principal component representation of mutational profiles, excluding PC1 because it correlates strongly with mean ER. Permissive positions have low ER scores for all substitutions and so the balance of other principal components is noisy. To account for this we split positions where the magnitude of all ER scores is less than 0.4 into a permissive subtype for each amino acid. Finally we use hierarchical clustering and hybrid dynamic tree cutting (Langfelder et al., 2008) to identify subtypes (**table S3**), which we label XP, XO, X1, X2, … for permissive positions, outliers and the main subtypes (in size order) of amino acid X. This approach gave us subtypes with distinct mean mutational profile patterns and therefore different properties, according to our exploration of the mutational landscape. 31 of 66 positions covered by multiple studies were assigned to the same subtype, and many others were in two functionally similar subtypes, despite differences in experiments (**figure S6**). This result is extremely unlikely by chance (one-tailed binomial test versus random assignment: *p* = 3.397 × 10^−4^) which suggests the procedure is robust.

We find between 1 and 8 subtypes for each amino acid, along with permissive positions for all 20 and outlier positions not consistent with any subtype for all but glutamine (**figure 3B**). In total this defines 100 amino-acid subtypes, including permissive positions for each amino acid. Histidine is the one amino acid where we only found a single subtypes, suggesting it tends to fulfil a similar role in most positions. A larger dataset would likely lead to discovery of rarer forms. The frequency of each successive subtype (**figure S7**) decreases exponentially across all amino acids (*f* = *e*^−0.56197±0.01655×*N*^, *r*^2^ = 0.9359, *p* < 2.2 × 10^−16^), suggesting that increasing the size of the dataset would have diminishing returns in terms of identifying new subtypes. It also suggests that the number of groups common enough to be considered a subtype is finite, with amino acids main functions covered well by this data and a potentially large number of rarer roles requiring a larger dataset to be discovered with these methods.

We found a number of common subtypes across amino acids, with similar mean mutational profiles (**figure 3C**). These subtype groups generally tolerate or select against a specific type of amino acid, for example requiring small aliphatic amino acids or tolerating everything apart from proline (referred to as the small aliphatic group and not proline group respectively). Subtype groups occupy different regions of UMAP space, thus tending to have different physical properties as well as different mutational profiles (**figure 3D**). The strongest is the ‘not proline’ group, whose positions’ only role appears to be forming the correct backbone conformation. These subtypes have a very consistent profile and are found in 14 of the 19 non-proline amino acids, occurring in up to 20.5% of amino acid positions. Methionine (20.5%) and glutamine (20%) have the most ‘not proline’ positions and larger amino acids, such as aromatics (mean 2.9%) or leucine (4.6%) tend to have fewer (**figure 3E**). They are often observed to occur in and around secondary structural elements, in particular having a significantly higher Porter5 likelihood (Torrisi et al., 2018) of occurring in alpha helices (mean 1.24 times as likely, one tailed Mann-Whitney U-Test: *p* = 1.7 × 10^−5^). Another notable group of subtypes only tolerate small aliphatic amino acids. Cross-referencing their positions in UMAP space with our property map (**figure 2**) reveals these positions tend to be highly conserved, buried, moderately hydrophobic and lead to Van der Waals clashes when mutated; which is exactly what would be expected of positions that specifically require small amino acids. The other subtype groups identified follow similar patterns, tolerating certain specific groups of amino acids (aliphatic, negative, not aromatic etc.). These subtype groups capture the major divisions of classic amino acid chemistry but, importantly, not all subtypes of each amino acid fall into the group matching their classical chemistry. For example, Y1 and Y3 are in the aromatic subtype group but Y2 tolerates hydrophobicity more broadly, and R1 selects for positive charge but R2 positions primarily select against negative charge rather than requiring the positive charge itself.

The most and least permissive subtypes of each amino acid vary widely in frequency (**figure 3E**), indicating how often each amino acid fills either highly specific or very general roles. For example, the most selective subtype of cysteine, methionine and tryptophan all occur in at least 65% of their positions in our data, meaning these amino acids are very often used for specific functions. Conversely, a much smaller proportion of glycine positions are the most selective subtype (27%) but that subtypes’ mean ER profile (**figure 3F**) shows those positions are some of the most selective in the dataset; it’s not common for a glycine to be highly selected but when they are they fulfil a very specific role.

The frequency of permissive positions varies between amino acids as well. Hydrophobic amino acid positions (e.g. aliphatic and aromatic), tend to be permissive significantly less frequently than other amino acids (mean 0.42 times as likely, one tailed t-test: *p* = 0.0001264), with this effect being particularly strong for aromatic amino acids (all <4% frequency). This may be because they tend to occur in positions that at least require hydrophobicity, such as the core of the protein, and therefore at least strongly hydrophobic substitutions are selected against. Permissive subtype positions also tend to be more surface accessible in general (mean 28.9Å^2^ greater accessible surface area, one tailed t-test *p* < 2.2 × 10^−16^, Cohen’s *d* = 0.61). The fact that hydrophobic amino acids are often experimentally tolerated on the surface but selected against in nature suggests the experiments may be missing some natural selection pressures.

### Characterising amino acid subtypes

The results so far show that broad subtype analysis can shed light on general trends in amino acid roles, but more targeted analysis is required to understand subtypes specific properties. We have chosen three groups of subtypes to cover in detail. Firstly, cysteine positions are divided between two main subtypes. The larger subtype, C1 (65% of positions), is generally intolerant to substitutions with aromatics being the most tolerated, whereas C2 (26.3%) tolerates any hydrophobic residue (**figure 4A**). C1 positions tend to be involved in cysteine specific functions, in particular disulphide bonds (**figure 4B**) and various ligand binding interactions (**figure 4C**). Conversely the majority of C2 positions are buried (**figure 4D**) and primarily utilise hydrophobic properties. Both subtypes also appear to be involved in interactions with aromatic residues via their π-orbitals (Orabi and English, 2016) (**figure 4E**).

**Figure 4.**
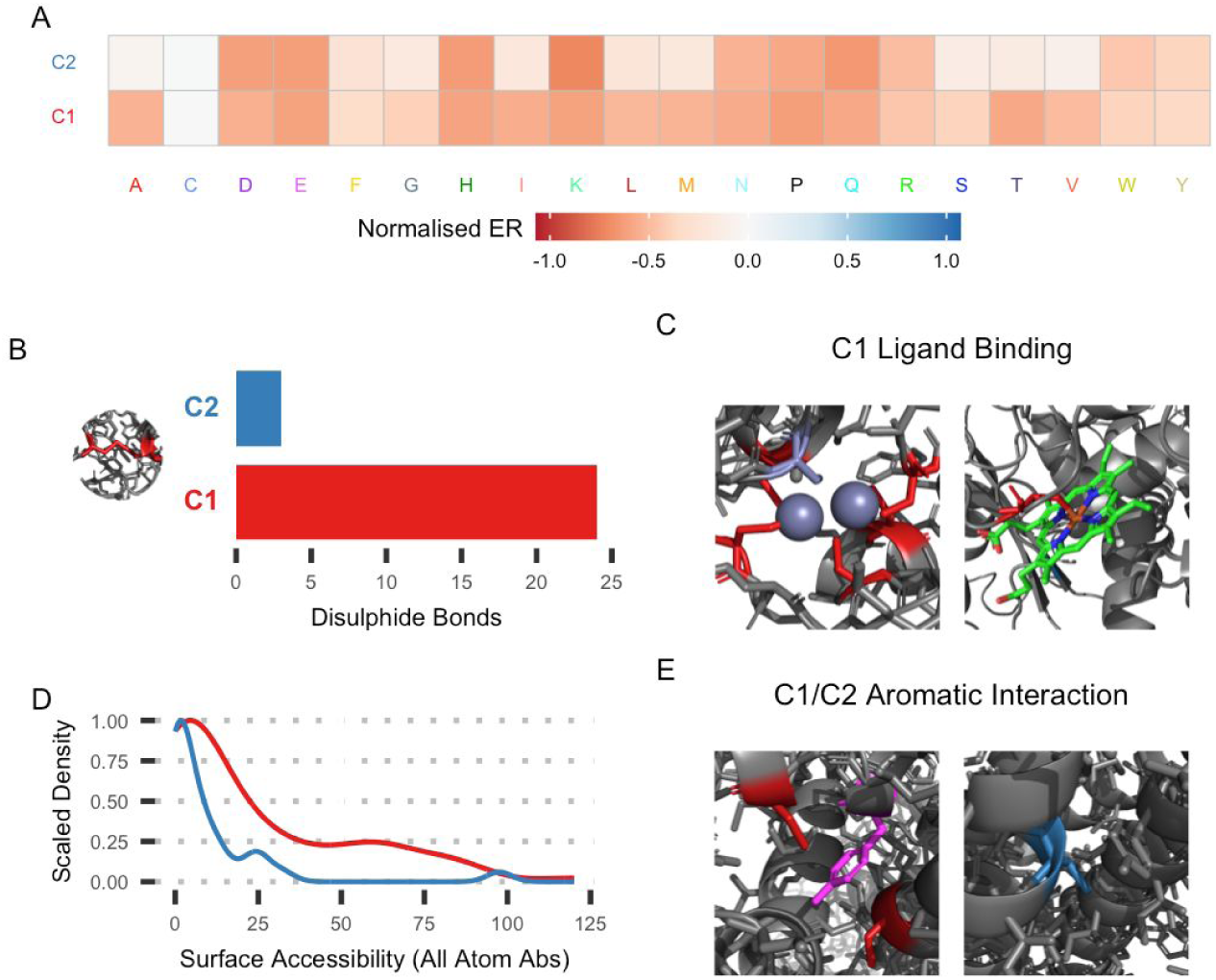
Cysteine subtype examples. **A:** Mutational profiles of the two cysteine subtypes. **B:** Number of positions of each subtype in a disulphide bond, based on FoldX prediction. **C:** Examples of C1 positions involved in zinc ion (left, GAL4, PDB ID: 3COQ) and haem (right, CBS, PDB ID: 4L0D) ligand binding. **D:** All atom absolute surface accessibility of cysteine subtypes. **E:** Examples of C1 (left, NP, homology model on PDB ID: 2Q06) and C2 (right, CCR5, PDB ID: 6MET) interacting with aromatic groups.

Another interesting set of subtype groups is those of the charged amino acids; negatively charged aspartate and glutamate and positive arginine and lysine. These visually split into 12 subtypes that fall into 5 broad categories (**figure 5A**), with the frequency of charged amino acid positions falling in each category noted: selective for each of the two charge polarities (26.7% negative and 21.1% positive); selective for general polarity (7.83%); selection against negative charge (6.18%) and selection against proline only (12.2%), leaving 26.1% in rarer subtypes, permissive positions and outliers. These groups are quantitatively differentiated by the average electrostatic force they contribute to the protein (**figure 5B**). The two groups that require charge tend to have the strongest forces, then the polar subtype group and the two selecting against properties have the weakest. A number of examples illustrate the typical roles of these different subtypes. Positions in the polar subtype group frequently interface with solvent (**figure 5C**). Positive and negative positions together form ionic bonds (**figure 5D**) and independently bind substrates such as DNA (**figure 5E**) or ions (**figure 5F**). Positions selecting against negative charge are often near negative charges that would repel them, either other residues in the protein or bound substrates such as RNA (**figure 5G**) and, as is generally typical, positions selecting against proline occur where fold topology is important, particularly at the ends of secondary structures (**figure 5H**) or in loops connecting them. The division of roles between subtypes is not absolute but the majority of cases of each role examined in structural models fall into the appropriate subtype, which is also the case with other examples mentioned (see **table S4**).

**Figure 5.**
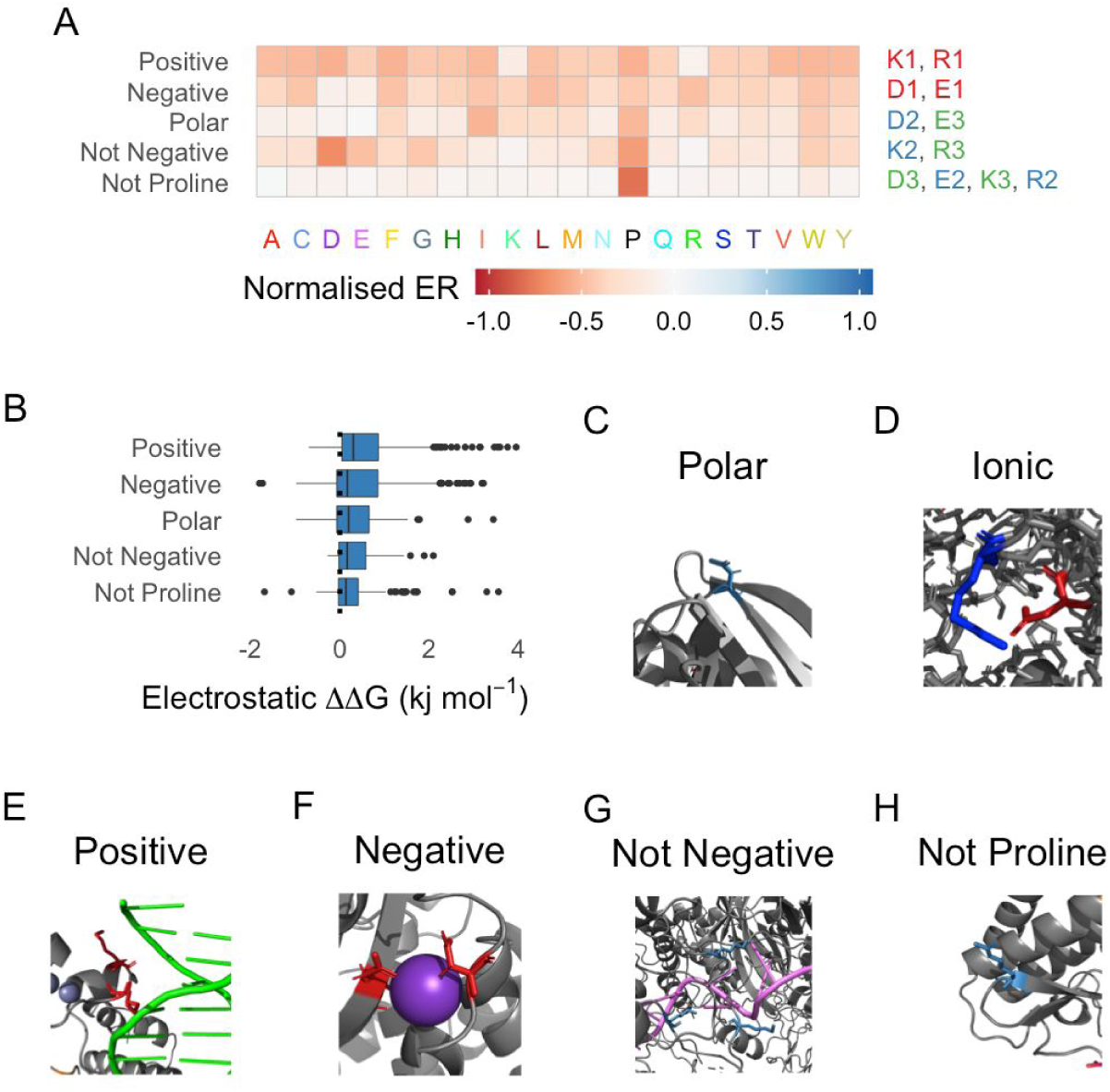
Polar subtype examples. **A:** Mutational profiles of polar subtypes **B:** Boxplots showing the distribution of the mean electrostatic component of FoldX substitution ΔΔ*G* at each position of the subtype group. **C:** Polar position on the protein surface (TEM1, PDB ID: 1M40). **D:** Example ionic interaction between positive and negative subtype positions (CBS, PDB ID: 4L0D). **E:** Positive subtype position interacting with the DNA backbone (GAL4, PDB ID: 3COQ). **F:** Negative subtypes binding an ion ligand (TEM1, PDB ID: 3COQ). **G: ‘**Not negative’ subtype positions surrounding bound RNA (PAB1, PDB ID: 6R5K). **H: ‘**Not proline’ position at the end of an alpha helix (APH3II, PDB ID: 1ND4).

The final example covers groups of subtypes requiring different sized hydrophobic amino acids, with one group of subtypes selecting for aromatics (F1, W1, Y1), one larger aliphatic amino acids (I2, L2, M1) and one small amino acids (A1, G2, P3) (**figure 6A**). The frequency that positions of each amino acid fall into these groups varies a lot. For example, 36.5% of phenylalanine positions and 34.8% of tyrosine positions are in the ‘aromatic’ group compared with 67.8% of tryptophan positions, suggesting tryptophan positions more commonly primarily utilise aromatic characteristics. The typical roles these subtypes have are illustrated by the average structural consequences of mutations at their positions, based on components of FoldX’s substitution ΔΔ*G* predictions. Firstly, subtypes requiring small amino acids make a bigger contribution to entropy minimisation during folding, having fewer possible configurations and helping the protein fold efficiently (**figure 6B**). They also frequently occur in cramped spaces within the proteins, resulting in clashes when mutated (**figure 6C**). On the other hand, larger amino acids, both aromatic and aliphatic, contribute more Van der Waals forces (**figure 6D**), highlighting the trade-off between Van der Waals forces, packing the correct folds and entropy minimisation when selecting for amino acids in the hydrophobic core. Finally, aromatic positions are more likely to be surface accessible than either aliphatic position (**figure 6E**, mean 16.8Å2 greater accessible surface area, one tailed t-test: *p* = 8.909 × 10^−6^), including when the slightly polar tyrosine is excluded. This potentially occurs because the aromatic ring interacts somewhat with water through its delocalised π-orbitals (Slipchenko and Gordon, 2009). Together, these examples illustrate some properties and typical roles of subtypes and show that the subtypes have biological and structural features in specific cases as well as in bulk statistics. In total, we have found 100 subtypes, characterised in **figures S8-27** and described qualitatively with examples in **table S4.**

**Figure 6.**
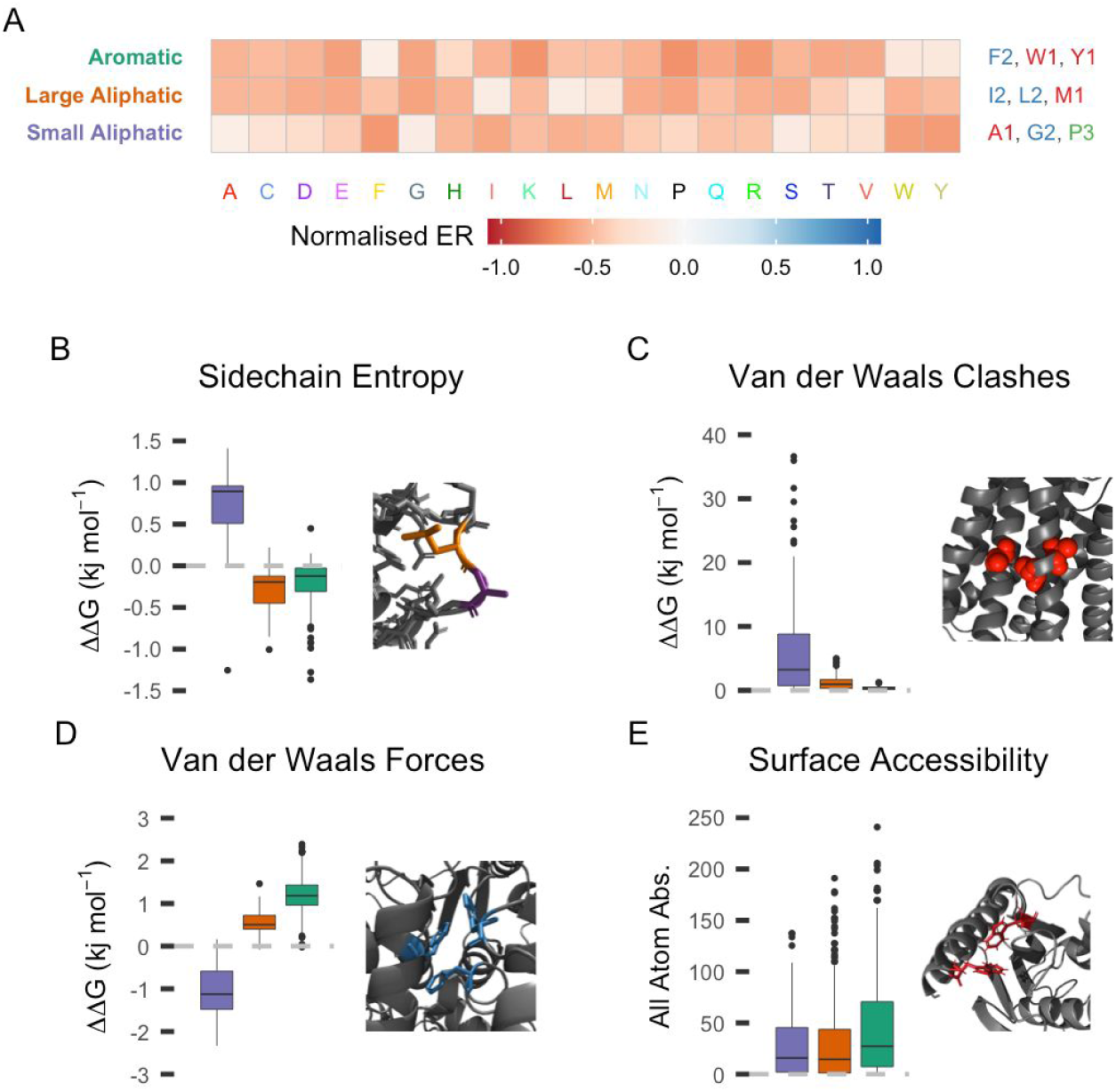
Different sized aliphatic subtypes. **A:** Mutational profiles of size conscious aliphatic subtypes. **B, C, D:** Boxplots showing the distribution of mean FoldX substitution ΔΔ*G* across positions of each subtype group, for various FoldX terms. **B:** Sidechain entropy and an example showing why mutations at small aliphatic group positions increase entropy more. **C:** Van der Waals clashes and an example showing alanine positions in a helix bundle demonstrating why clashes occur on mutation of small residues. **D:** Van der Waals forces and an example showing internal aromatic groups making Van der Waals interactions. **E:** Boxplot showing the distribution of all atom absolute surface accessibility between groups and an example of surface aromatic groups.

## Discussion

We have demonstrated the utility of combining deep mutational scanning studies for performing proteome scale analyses, even when the experiments differ between studies. While the results of these studies are strongly related to evolutionary conservation, they also provide quantitative information about rare variants that would require prohibitively large sample sizes to measure through sequencing. As we demonstrate here, this makes them useful for analysing mutational profiles because these frequently include rare variants.

The mutational landscape we describe clearly demonstrates the link between mutational profiles and biophysical properties. It is a potentially powerful tool for data driven characterisation of a position’s likely properties using mutational profiles or conversely estimation of mutational profiles from properties. This can be thought of as a continuous form of subtypes, where each position is described by a vector of properties, such as the principle components or UMAP transformation of its mutational profiles, and these vectors indicate biophysical properties of the position. These representations could also be useful for predictive models, especially if the landscape was developed with more data and more sophisticated methods linking it to properties.

The discrete classification of positions into amino acid subtypes provides similar power in a clearer form, providing a mapping between types of position and mutational profiles, as well as providing unique benefits such as the ability to more clearly quantify the range of diverse functions each amino acid fulfils and their frequencies. In this work we identify 100 subtypes including permissive positions of each amino acid but excluding some outlier positions. In addition we find interesting differences in the frequency of these roles, for example cysteine positions seem to utilise cysteine specific properties in 65% of positions while charged residues only rely heavily on their charge in 21-27% of positions.

Subtype characterisation also breaks down typical roles of amino acids and their frequencies, which helps quantify the spread of functions across the proteome. The subtypes we identify include both amino acid specific functions like ion binding and general roles, such as selection against proline or for hydrophobicity. This highlights another way to view some subtypes; they are shared between amino acids, and so an alternative approach would be to combine these subtypes, either after subtypes have been generated or by clustering all amino acids together. Such an analysis could help further quantify amino acid diversity by explicitly identifying positions where several amino acids share the same properties.

The relationship between deep mutational scanning and evolutionary conservation suggests that a similar mutational landscape analysis could be performed using sequence alignment data. We experimented with applying a conservation based analysis to our dataset, based on positions SIFT score profile. The SIFT based mutational landscape captures many of the same properties as that based on deep mutational scanning, although not the positioning of transmembrane protein domains (**figure S28**). However, subtypes produced using the same algorithm on log_10_SIFT score profiles fail to capture many of the roles that ER base subtypes do. For example, disulphide bonds are shared evenly between the two SIFT based cysteine subtypes, ‘not proline’ type subtypes are not identified and aspartate positions binding ligands are not separated as clearly. In addition, ER based subtypes average profiles are more differentiated from other subtypes of the same amino acid than SIFT based subtypes (**figure S29**, mean 0.13 greater cosine distance, one-tailed Mann-Whitney U-Test: *p* < 2.2 × 10^−16^). This suggests that conservation based subtypes are a potentially powerful future direction, allowing coverage of a much larger portion of the proteome, but lack the resolution and detail provided by deep mutational scanning approaches. A method combining both data types could also be powerful.

Exploring how to combine and use this type of data will be important in future because many more deep mutational scanning experiments are expected. The most recent DepMap release (Dempster et al., 2019; DepMap, Broad, 2020; Meyers et al., 2017) suggests 7293 genes are essential in at least one cell line, and thus directly targetable by deep mutational scanning. This suggests data from 1000s of genes could be available in future, ideally from more standardised experiments. While we have data from a diverse range of proteins and have amassed as large a dataset as possible, we are ultimately limited to data from 6291 positions; a very small slice of the proteome. This is part of the reason we do not, and did not expect to, identify clear novel amino acid functions from our data; we only had access to data from a limited range of well studied proteins. Potentially a larger range of proteins would contain enough examples of rare, unstudied interactions to identify novel roles.

A larger dataset has potential both for extending this work and exploring other areas, for example a proteome wide analysis of different types of substitution in different contexts (i.e. focussing on the mutant amino acid rather than the wild type). For our work, a larger dataset has the potential to expand the range of roles discovered, with rarer subtypes either completely missing or too rare to identify in our data. The 100 amino acid subtypes need to be considered a lower bound estimate of amino acid roles in proteins that can be derived in this way. A form of subtype we would expect to find with more data is post-translational modification sites such as phosphosites. However, we currently only cover 52 phosphosites (The UniProt Consortium, 2019), and these are not all necessarily conserved, active sites. These sites would potentially tolerate other phosphorylatable residues and phosphomimetic amino acids. Another potential outcome is dividing subtypes we have identified into more specific forms, for example splitting C1 into disulphide bonding and ligand binding positions. Thus, our estimates form lower bounds for subtype numbers, although with diminishing returns as you add more data. Finally, all the subtypes we identify are spread between at least 7 fairly dissimilar genes, suggesting we are not identifying roles specific to classes of protein, for example in active sites, which could be discovered with more data giving more examples of such specific functions.

Overall, our analysis shows three key points. Firstly, deep mutational scanning data can be combined from disparate studies into a meaningful dataset that relates to real biology. Secondly, the mutational landscape is a powerful tool for analysing protein positions, with rich relationships to biophysical properties. And finally, positions of each amino acid can be broken down into typical subtypes using these profiles, allowing us to quantitatively map the diverse range of functions each amino acid plays across the proteome.

## Methods

The code to perform all the analyses in this study are available at https://github.com/allydunham/aa_subtypes. The combined dataset and subtype assignments are available as supplementary data files (**table S2** & **S3**).

### Combining the Data

First we selected as many deep mutational scanning studies (**table S1**) as we could find that fulfilled the following criteria:

- Available data
- Selection criteria matching the proteins natural function
- Scores that cannot be transformed to the standard 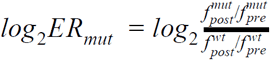 form, where 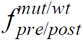 is the frequency of of variant *mut* or the *wt* before or after selection

This included searching MaveDB (Esposito et al., 2019) and searching the literature. Each study was then processed into a standard state, with a complete set of variant scores for single substitutions at each position.

Some studies generated multiple mutations to individual sequences, without generating all single substitutions as well. In these cases we used the average ER score of sequences including each substitution that wasn’t directly tested, limiting the average to sequences with a maximum number of variants, depending on the deviation from known single substitution scores and resultant variant coverage (**figure S1, table S1**). In studies with multiple comparable replicates, either direct replicates or in equivalent conditions, we averaged across conditions, and when multiple incomparable conditions were available we chose the most representative of proteins’ natural functions (**table S1**). Studies were then filtered if they had very poor correlation with SIFT scores, indicating unrealistic selection criteria or experimental oddities, or low coverage of substitutions at each position.

The scores for each study were transformed onto the standard scale, with a different transform required in each case (**table S1**), and normalised by dividing all scores by the median score of the lowest 10%. This threshold was chosen heuristically to encompass the typical ER scores of nonsense mutations. These scores were put together into a combined dataset, from which positions with <15 of the 20 missense and nonsense substitutions were filtered and remaining missing data was imputed using the median of that substitution type (A → C, A → D, …) across all normalised scores. Synonymous substitutions were imputed to be 0 if not measured (some studies include variant codons and so measure synonymous variants’ ER).

We identified the best structural model for each protein in SWISS-MODEL (Waterhouse et al., 2018), selecting higher resolution and coverage where possible, as well as favouring experimental models over homology models. The models used are detailed in **table S1**.

### Analysing the Mutational Landscape

The combined mutational landscape data was annotated with additional biophysical data from a number of tools:

- SIFT4G – substitution effect scores (with a custom patch to output SIFT scores to 4.d.p)
- FoldX – substitution ΔΔ*G* predictions
- naccess – surface accessibility measurement
- Porter5 – secondary structure predictions

The different physical components of the FoldX ΔΔ*G* predictions were averaged across all substitutions at a position in most analyses, giving a measure of the importance of different structural effects at that position. The UMAP (McInnes et al., 2018; Melville et al., 2020) and principal component dimensionality reductions were calculated and cross referenced with those factors.

### Identifying Amino Acid Subtypes

We tried a number of methods to cluster amino acid positions, using various algorithms, distance metrics and profile formultations. Our final algorithm, which performed by far the best, was as follows, applied independently to positions of each amino acid:

1. Separate permissive positions (|*ER*| < 0.4 for all substitutions)
2. Apply hierarchical clustering to the remaining positions, with average linkage and cosine distance to profiles consisting of PC2 to PC20 (calculated across the whole mutational landscape).
3. Use hybrid dynamic tree cutting (Langfelder et al., 2008) to assign positions to subtypes, using deepSplit = 0 or 1 depending on the amino acid.

We decided to use principal component space, with PC1 excluded, and cosine distance to remove the influence of mean ER score, which otherwise dominated clustering. Splitting positions by overall functional importance would also be an interesting analysis, but masks differentiation based on biophysical role, which is what we were interested in. Using PC2 to PC20 achieves this because PC1 captures the variation in mean ER. Using the cosine distance also helps because it measures the angle between the vectors formed by two profiles in multidimensional space, which is independent of their overall magnitude. However, using this metric means the distance between low magnitude positions is very noisy, since their ‘direction’ is largely random. To combat this we manually separate these positions before clustering.

The deepSplit parameter to the dynamic tree cutting algorithm determines how much to split the dendrogram. We vary this parameter between amino acids, increasing it to 1 for amino acids that appear to under-split using 0, based on profile consistency in the resultant subtypes. This is potentially partially required due to the amount of data we have, and more data could allow us to optimise this parameter across the dataset as a whole.

Subtypes were characterised using the averages of the various statistics and metrics we previously annotated the mutational landscape with, as well as their average ER score profiles.

## Supplementary Information

**Figure S1.**
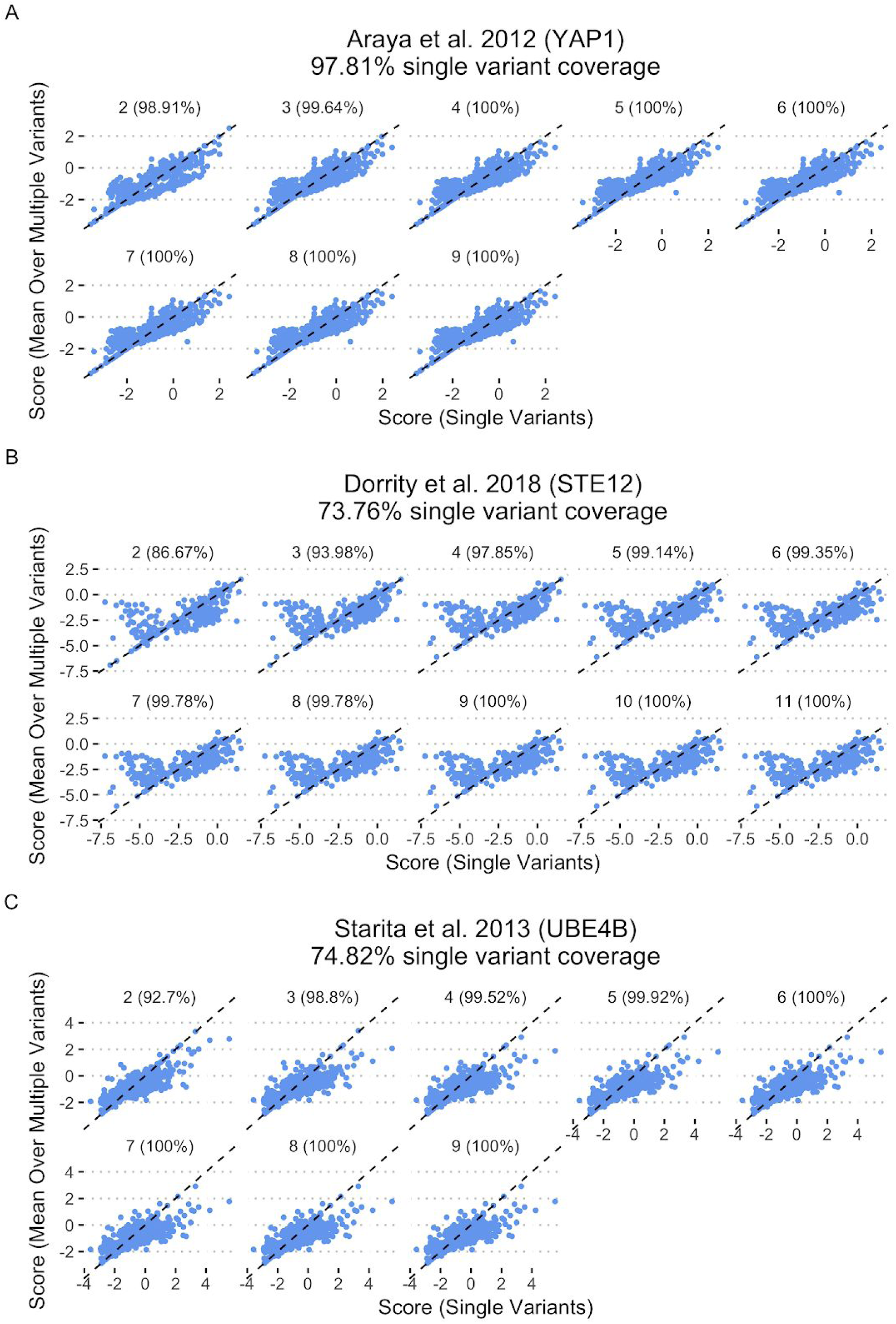
Validating averaging scores from multiply mutated sequences to estimate individual substitution scores in data from Araya et al. 2012 (**A**), Dorrity et al. 2018 (**B**) and Starita et al. 2013 (**C**). Each panel plots the score for explicitly measured single variants against the average score for multi-mutants containing that variant, using sequences with at most an increasing number of variants across subpanels. Each panel and subpanel also notes the score coverage of substitutions in the data when averaging sequences with that many mutations.

**Figure S2.**
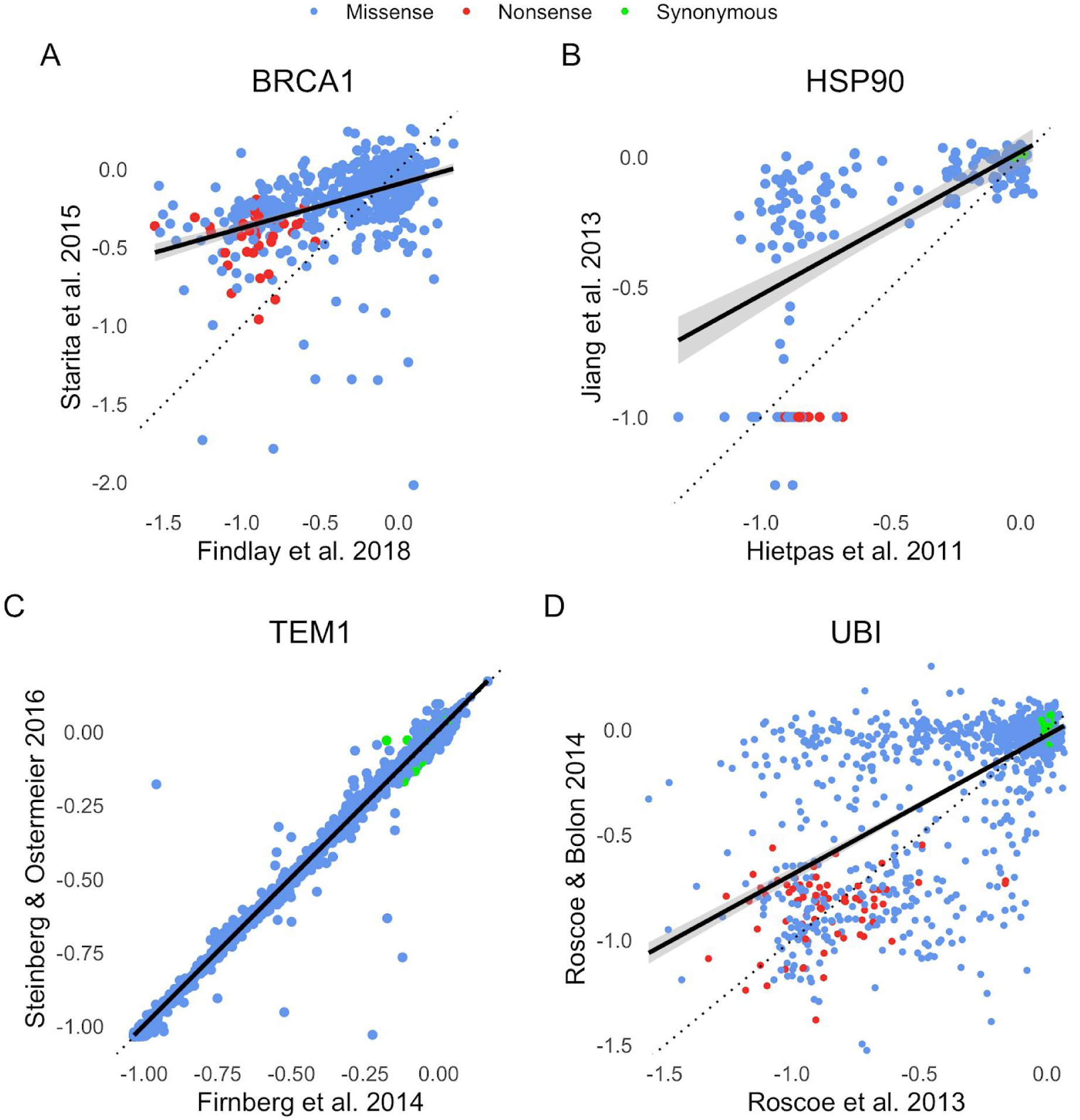
Correlation between scores from studies covering the same gene. **A**: BRCA1 (from the Fields and Shendure labs) **B**: HSP9 (both studies from the Bolon lab) **C**: TEM1 (both studies from the Ostermeier lab) **D**: Ubiquitin (both studies from the Bolon lab)

**Figure S3.**
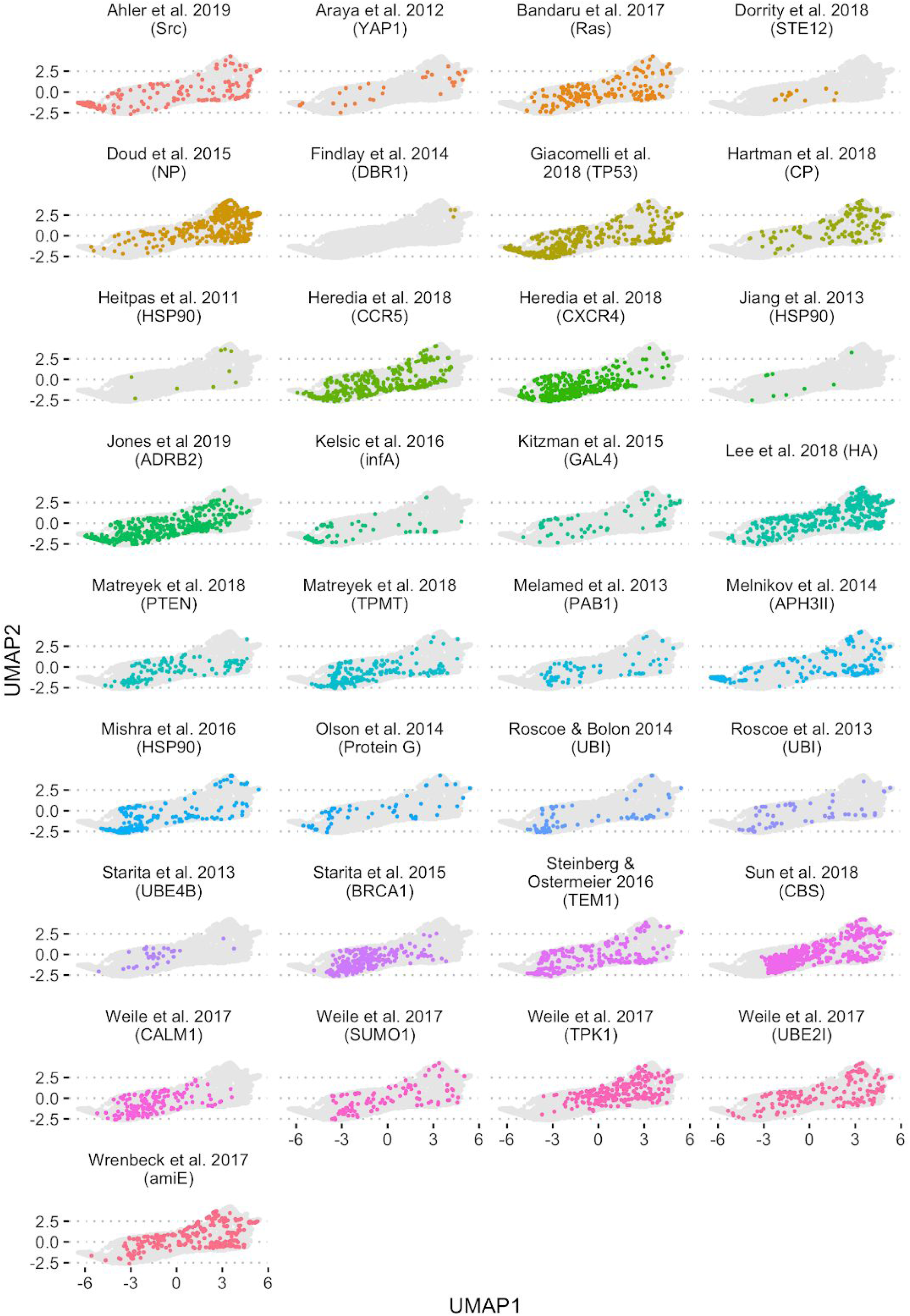
Positions from each study mapped to UMAP space. The positions from each study are coloured while the overall extent of the space is shown in grey, demonstrating points span the space rather than clustering by study. The few cases where some clustering appears are in studies with very few results, which are restricted to specifically chosen functional regions and so would be expected to have atypical landscape distributions.

**Figure S4.**
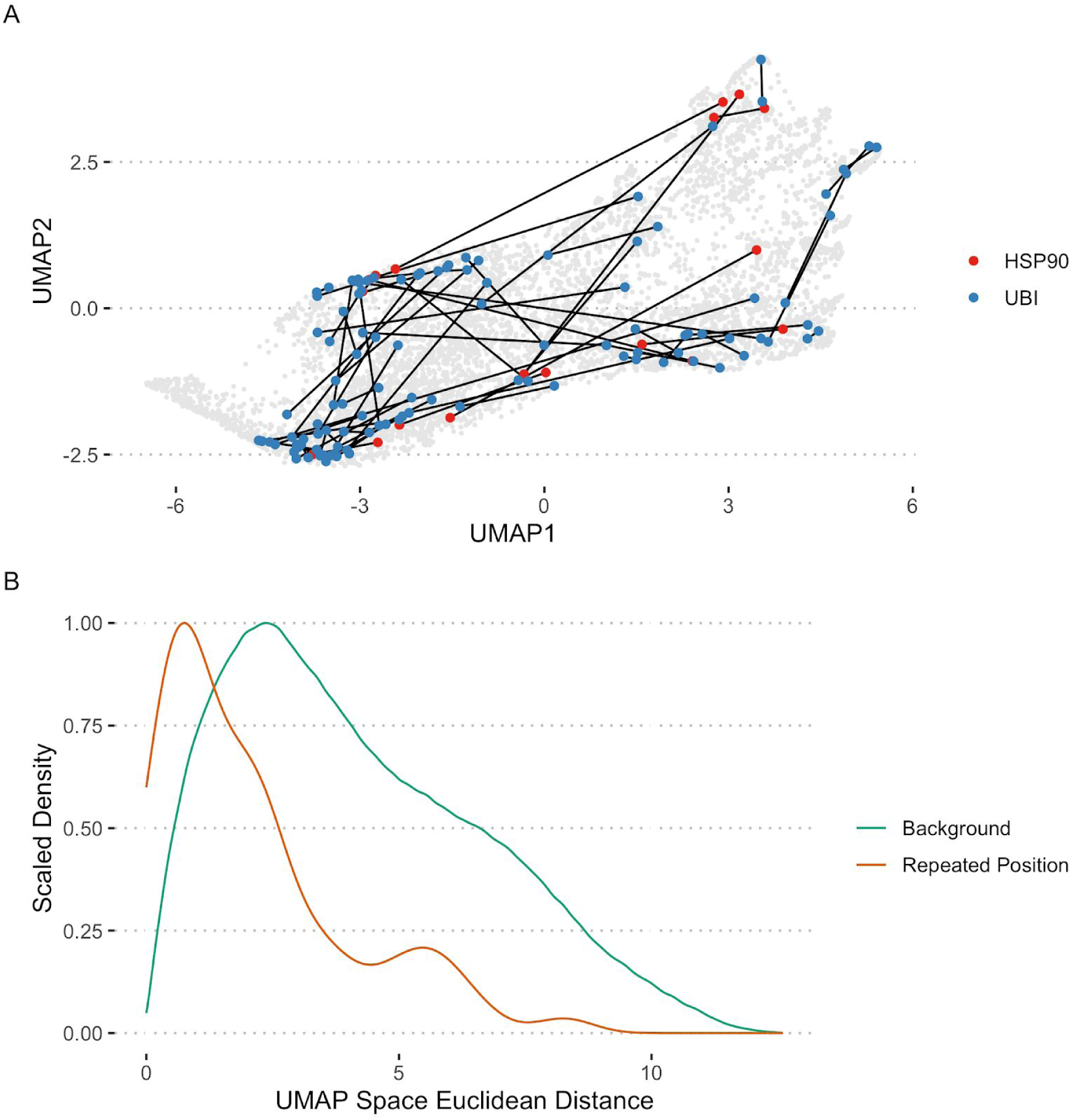
Distance between repeated positions in UMAP space. **A**: Repeated positions mapped into UMAP space, with the two copies of each position linked by a line. **B**: Distribution of distances between points in UMAP space, comparing repeated positions to the background distribution of random position pairs. Repeated positions are significantly closer on average (One tailed Mann-Whitney U-Test: *p* = 1.141 × 10^−13^).

**Figure S5.**
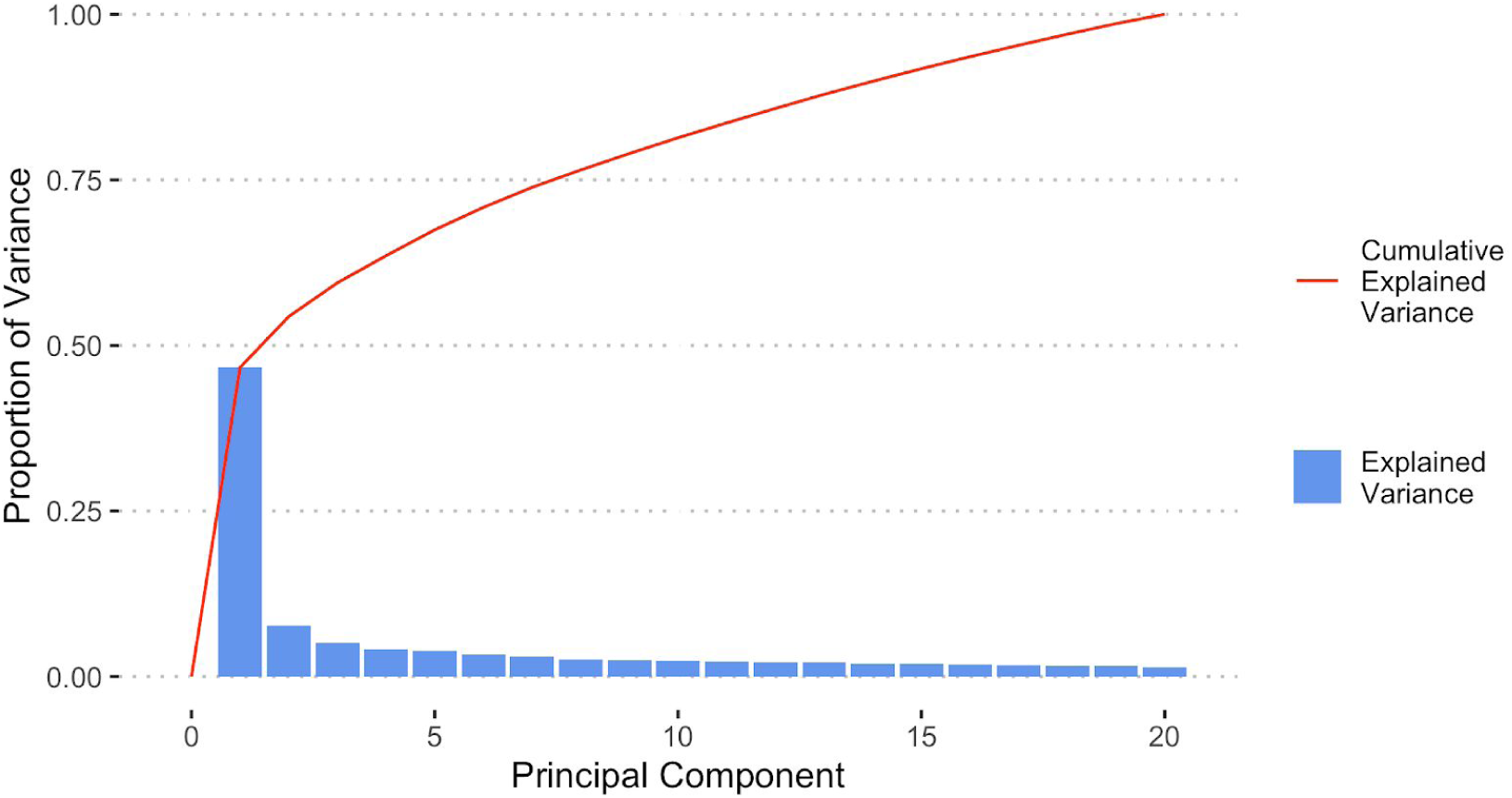
Proportion of variance explained by each principal component (blue bars) and the cumulative proportion explained (red line)

**Figure S6.**
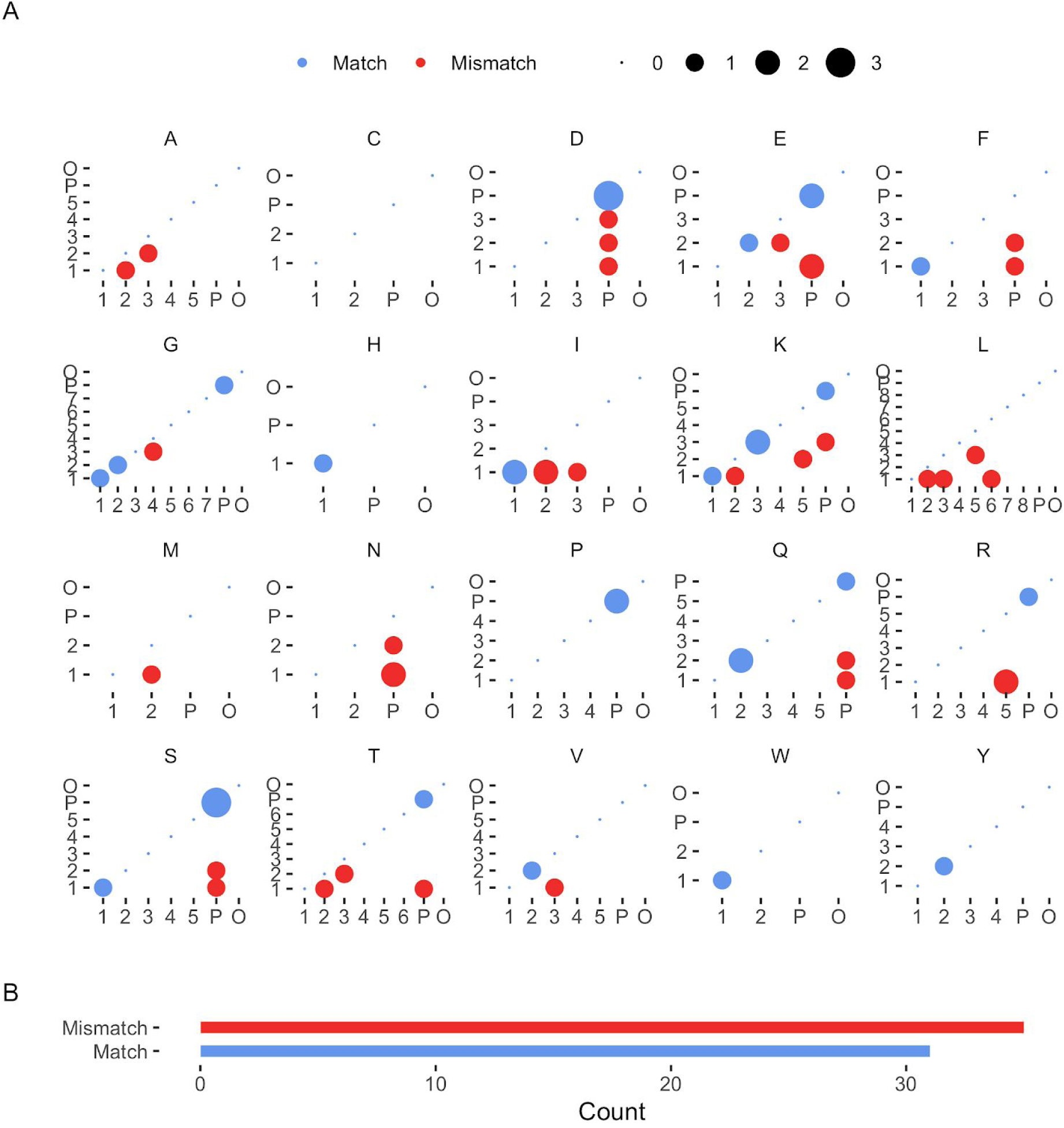
**A**: Pairs of subtypes the position covered by multiple studies were assigned to, showing matches in blue and mismatches in red. Subtype number pairs are sorted such that they are always placed in the lower triangle. **B**: Count of matches and mismatches.

**Figure S7.**
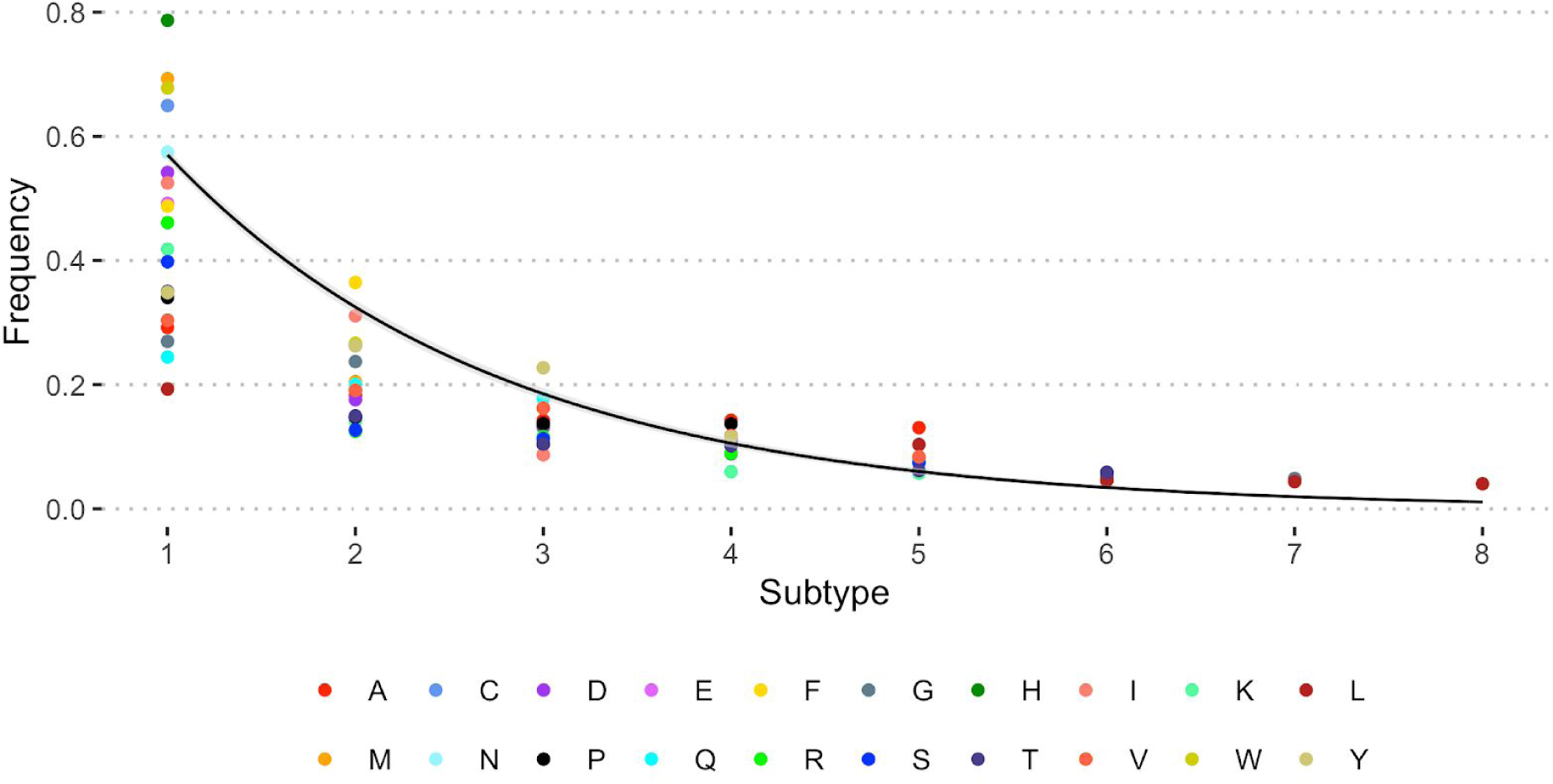
Frequency of each subtype number fit by an exponential function, across all amino acids. *y* = *e*^−(0.56197±0.01655)*x*^, *r*^2^ = 0.9359, *p* < 2.2 × 10^−16^

**Figure S8-27** – Plots characterising the subtypes of each amino acid, each following the same format. These are attached in an additional file. **A**: Number of positions assigned to each subtype. **B**: Relative probability of each subtype occurring in each Porter5 secondary structure **C**: ER profiles **D**: log_10_SIFT score profiles **E**: Surface accessibility distributions **F**: Normalised mean profiles of each FoldX energy term for substitutions at positions of each subtype **G**: Normalised mean chemical environment profiles for positions of each subtype. These comprised the count of each type of amino acid within 10Å of the target position.

**Figure S28.**
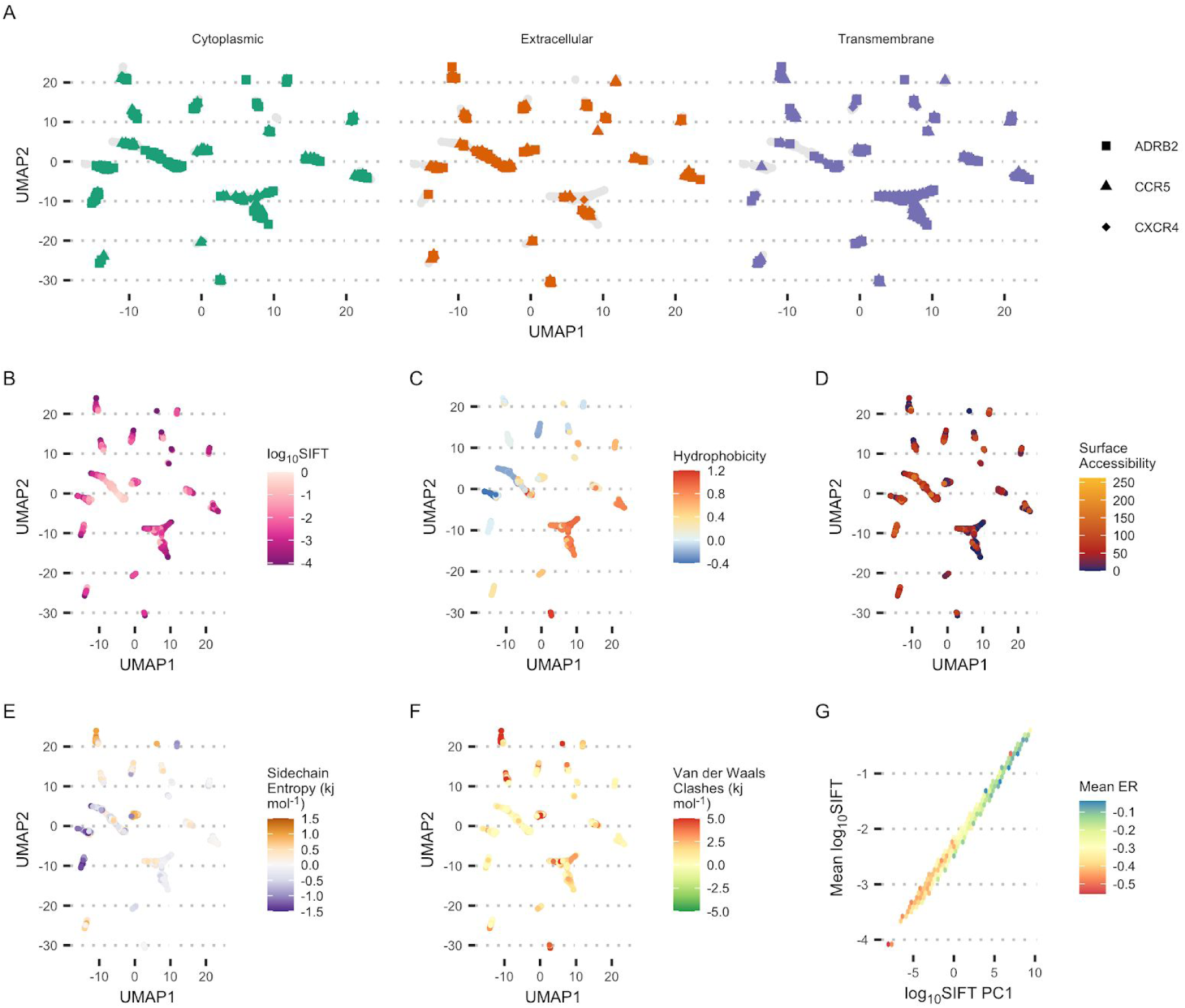
Mutational landscape based on SIFT score profiles (as a proxy for evolutionary conservation). **A**-**F**: Features mapped to UMAP space calculated from SIFT score profiles. **A**: domains of ADRB2, CCR5 and CXCR4. **B**: mean log_10_SIFT score. **C**: Average amino acid hydrophobicity. **D**: Surface accessibility **E**: Mean FoldX sidechain entropy substitutions at each position **F**: Mean FoldX Van der Waals clashes ΔΔ*G* term for ΔΔ*G* term for substitutions at each position. **G**: Correlation between PC1 (PCA on log_10_SIFT score profiles) mean log_10_SIFT score and Mean ER score.

**Figure S29.**
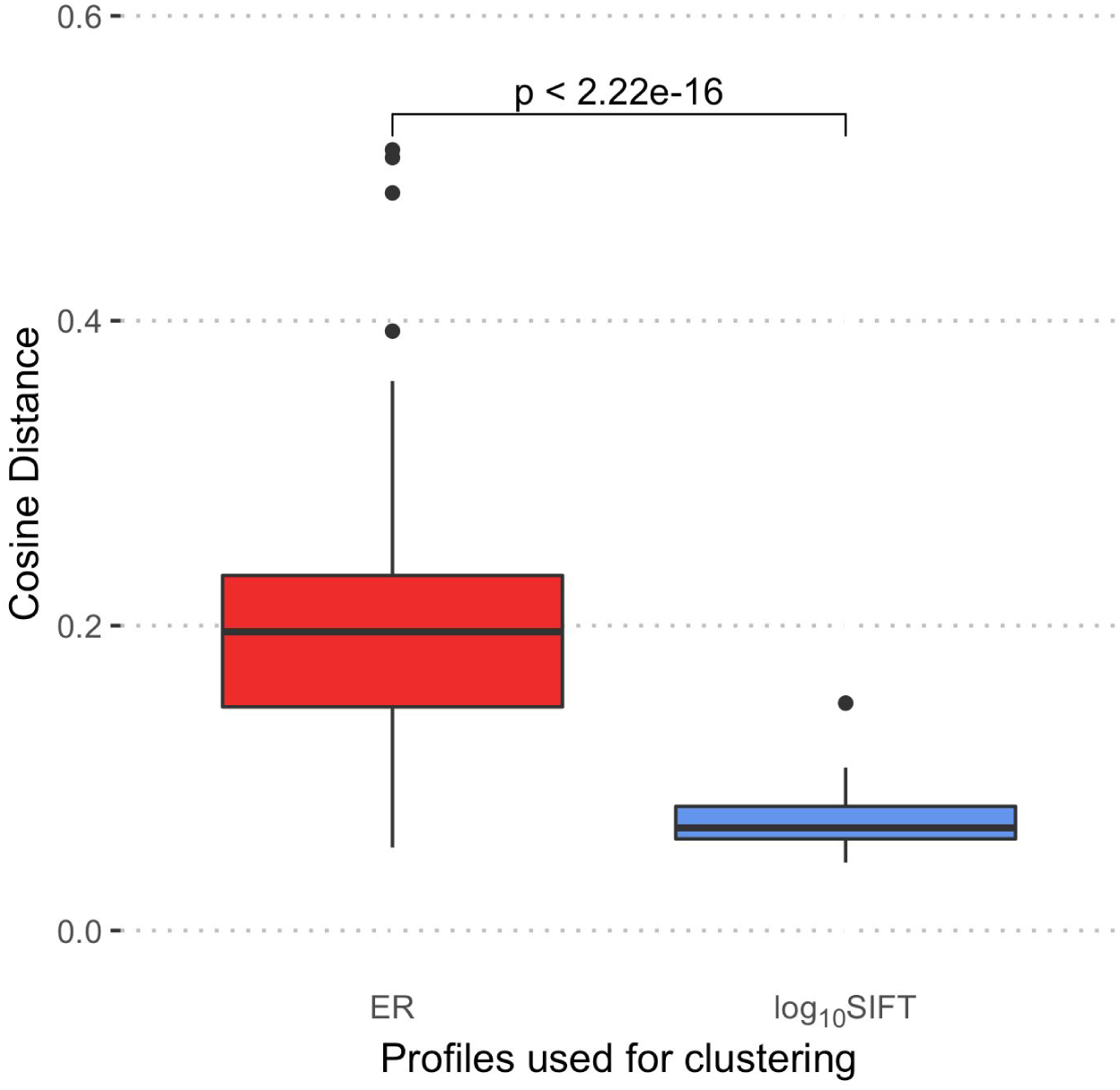
Comparison of average cosine distance from each subtype’s mean profile to other profiles of subtypes of the same amino acid. The profiles are ER scores for ER subtypes and log_10_SIFT scores for SIFT based clusters. The statistical test is a one tailed Mann-Whitney U-Test with the alternative hypothesis that ER subtypes are on average more different to each other.

### Supplementary Tables

**Table S1** – Studies selected for the paper, including those that were filtered after initial processing. Includes the number of positions/variants used, structural model chosen, transformation applied, condition and multiple variant processing and filtering notes.

**Table S2** – Deep mutational landscape. Lists all the positions used in the study along with associated data from SIFT, FoldX etc.

**Table S3** – Subtypes assigned to each position by our algorithm.

**Table S4** – Qualitative descriptions of each subtype (sheet1) and structural examples examined for this study (sheet2).

