## Supplementary figures and images for "Exploring amino acid functions in a deep mutational landscape"

### figureS8.pdf

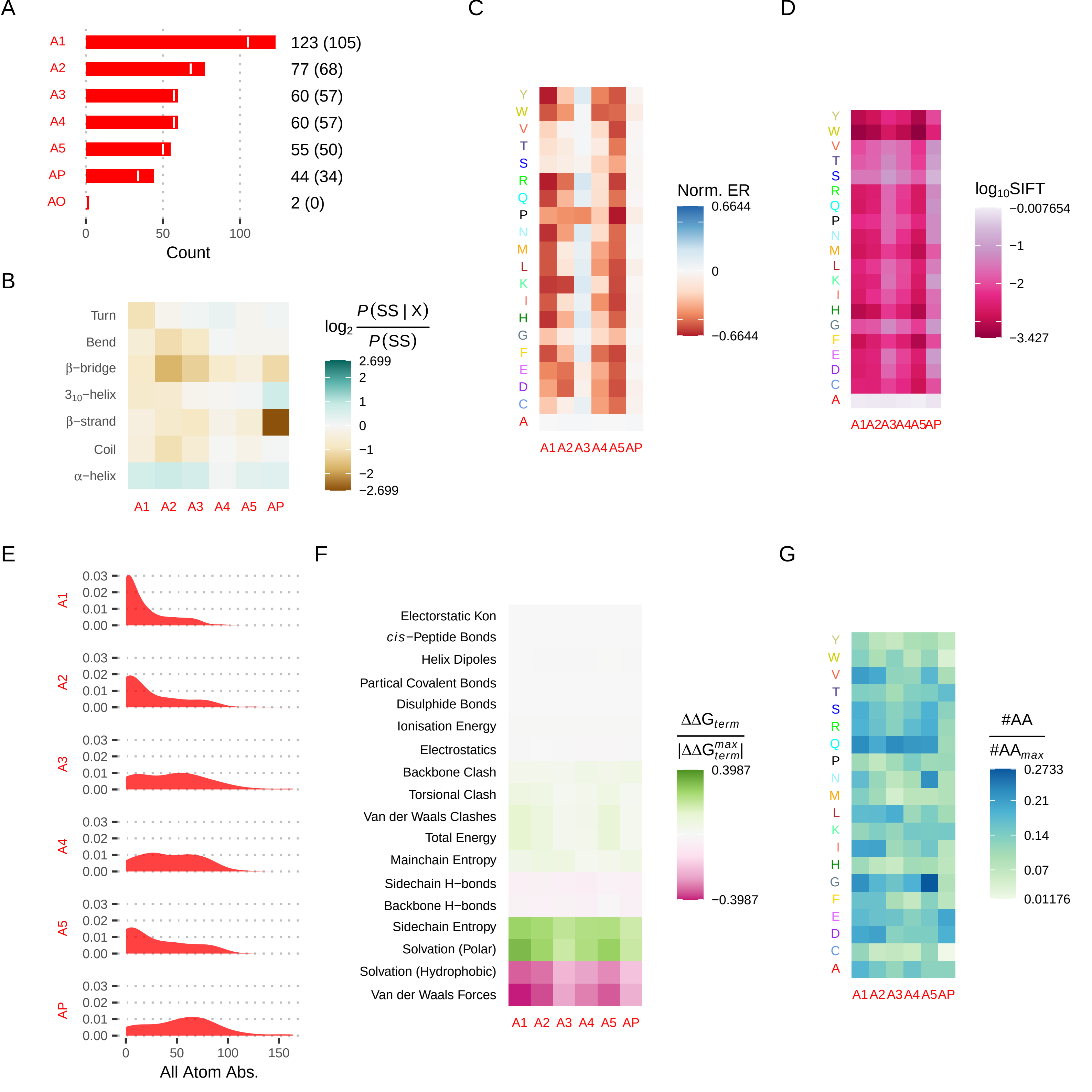

### figureS9.pdf

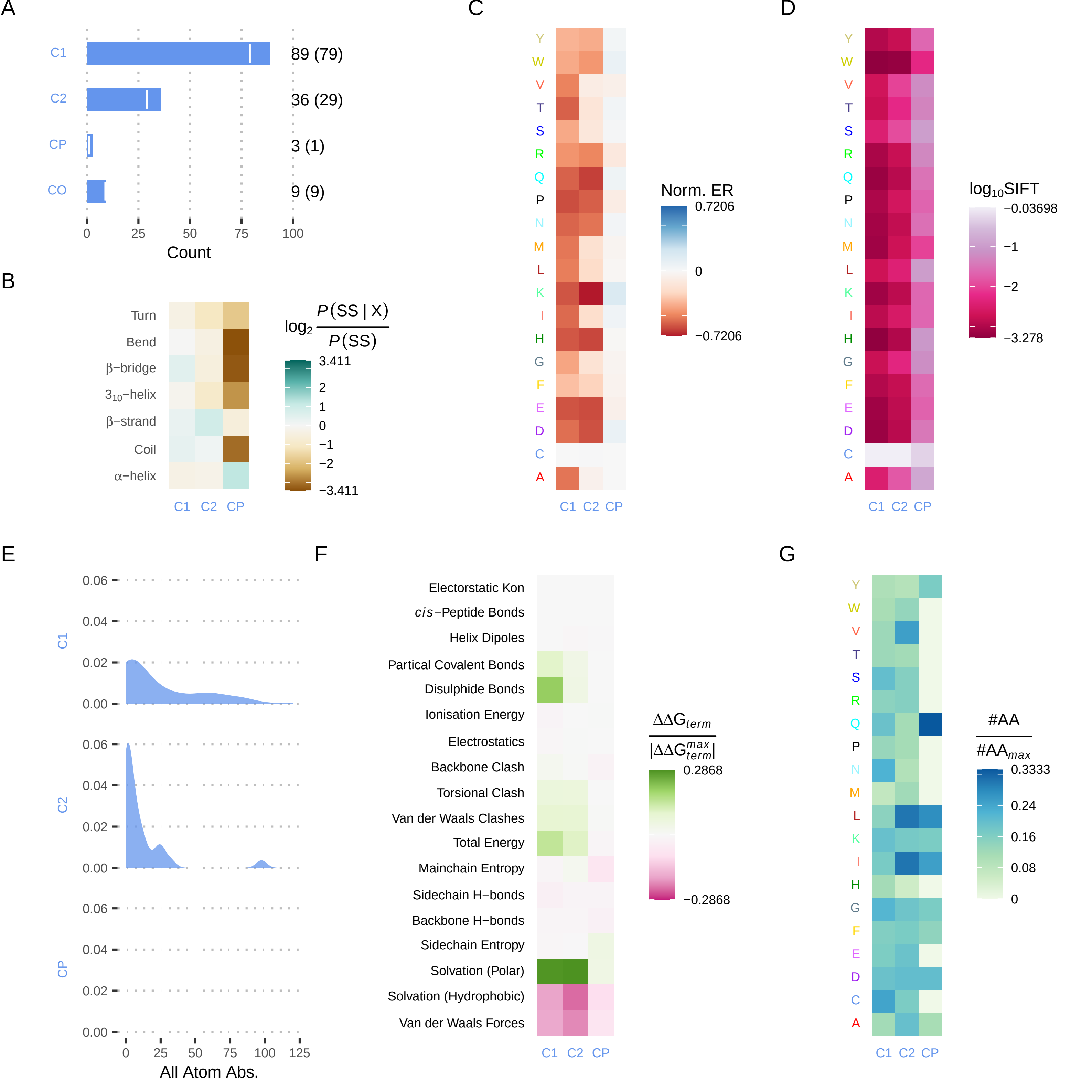

### figureS10.pdf

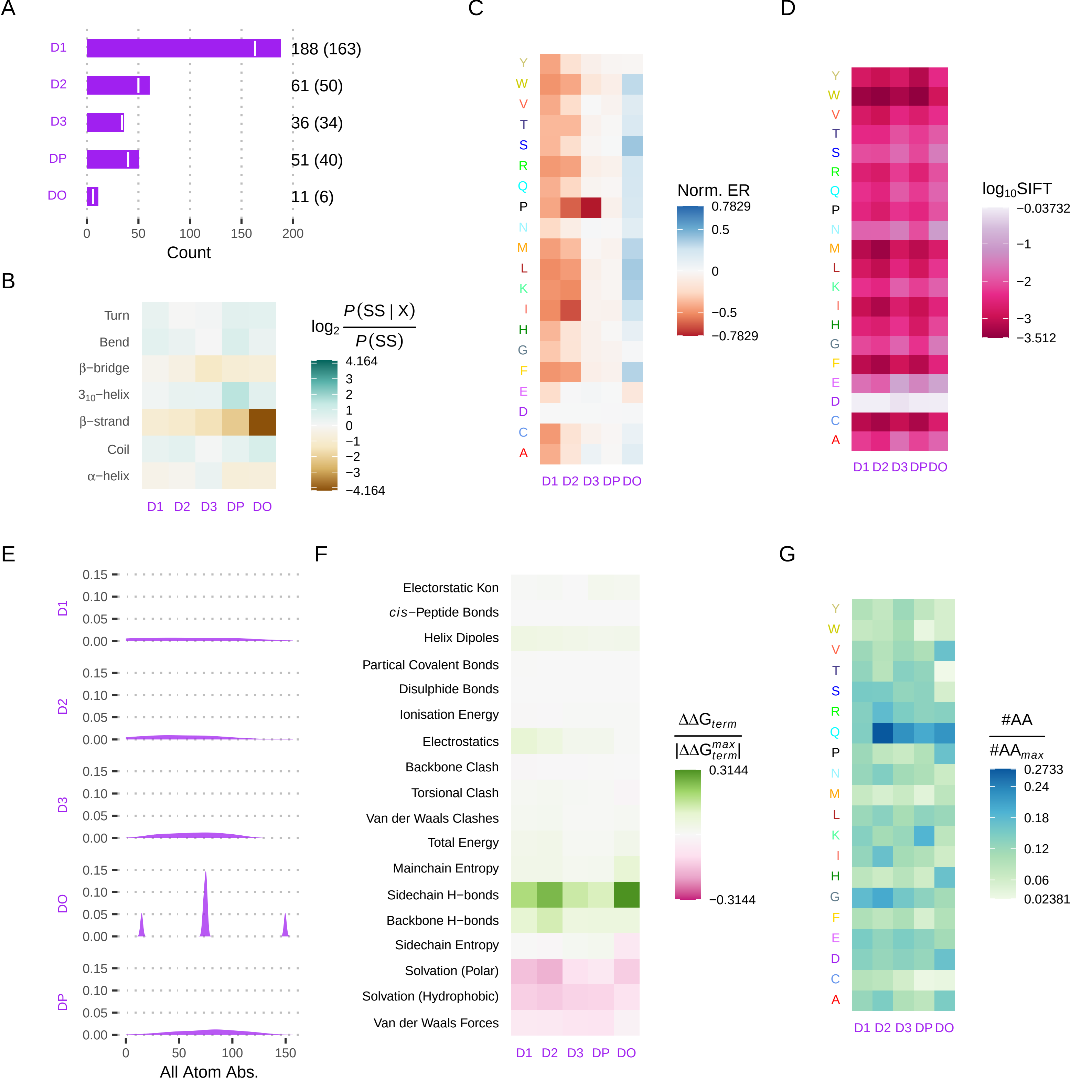

### figureS11.pdf

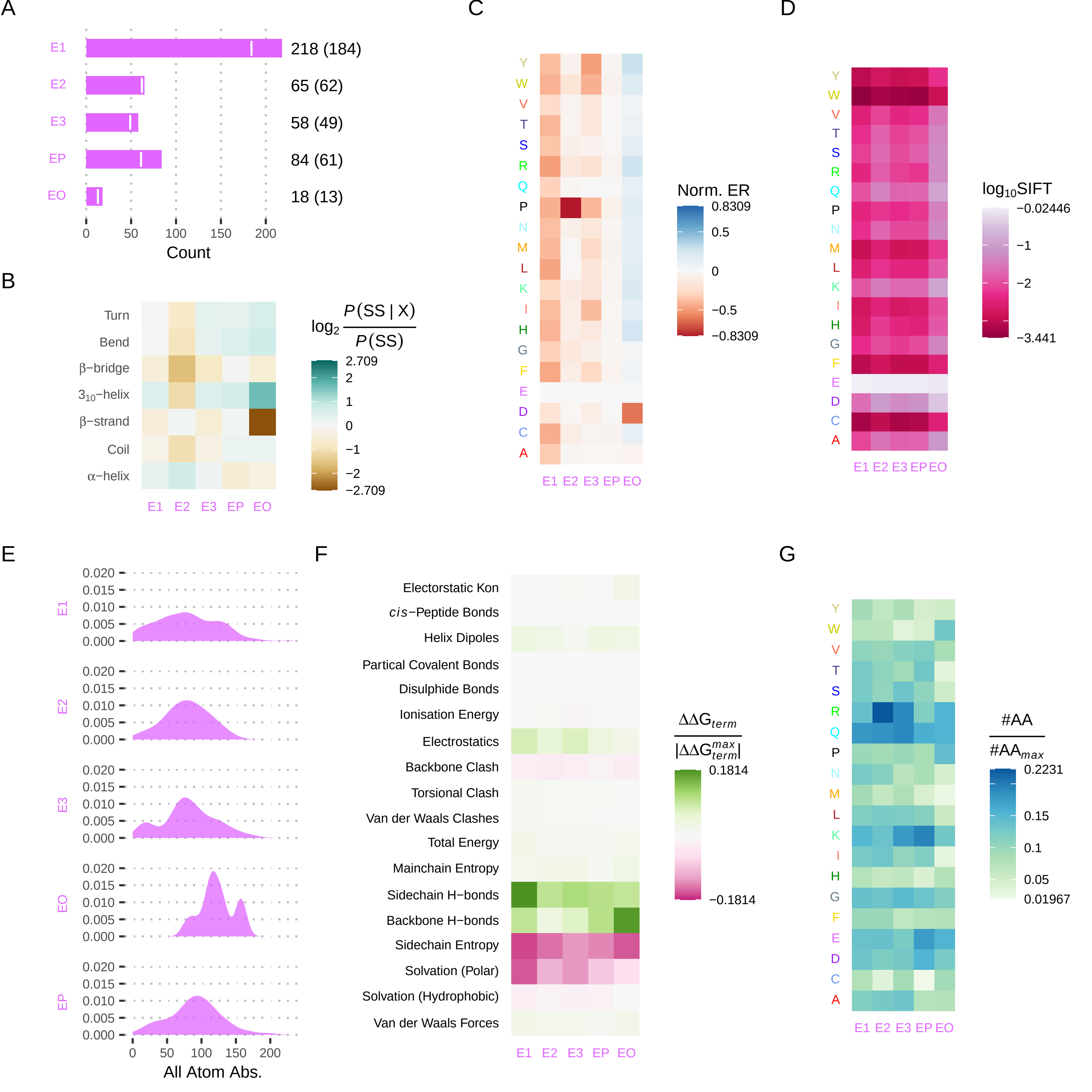

### figureS12.pdf

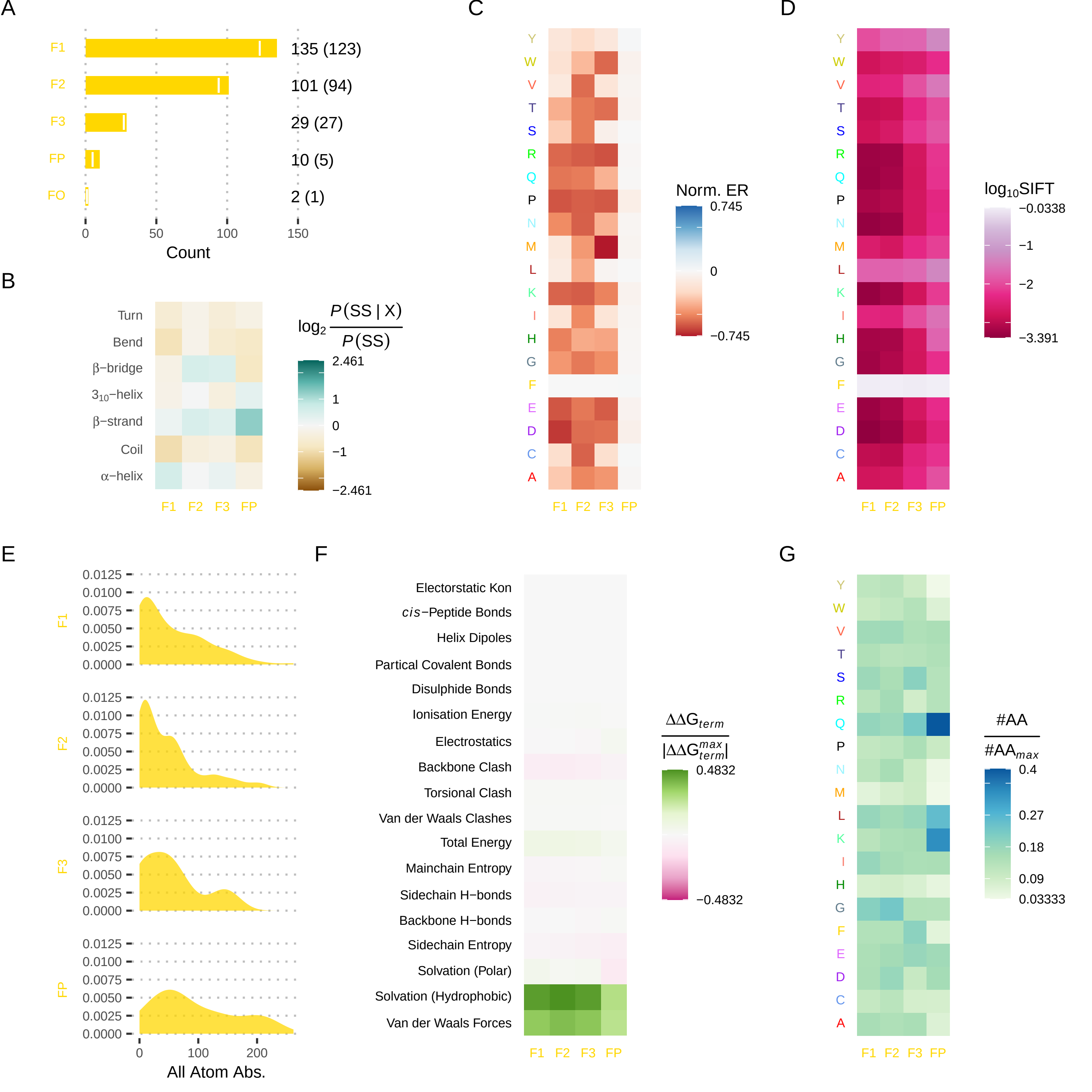

### figureS13.pdf

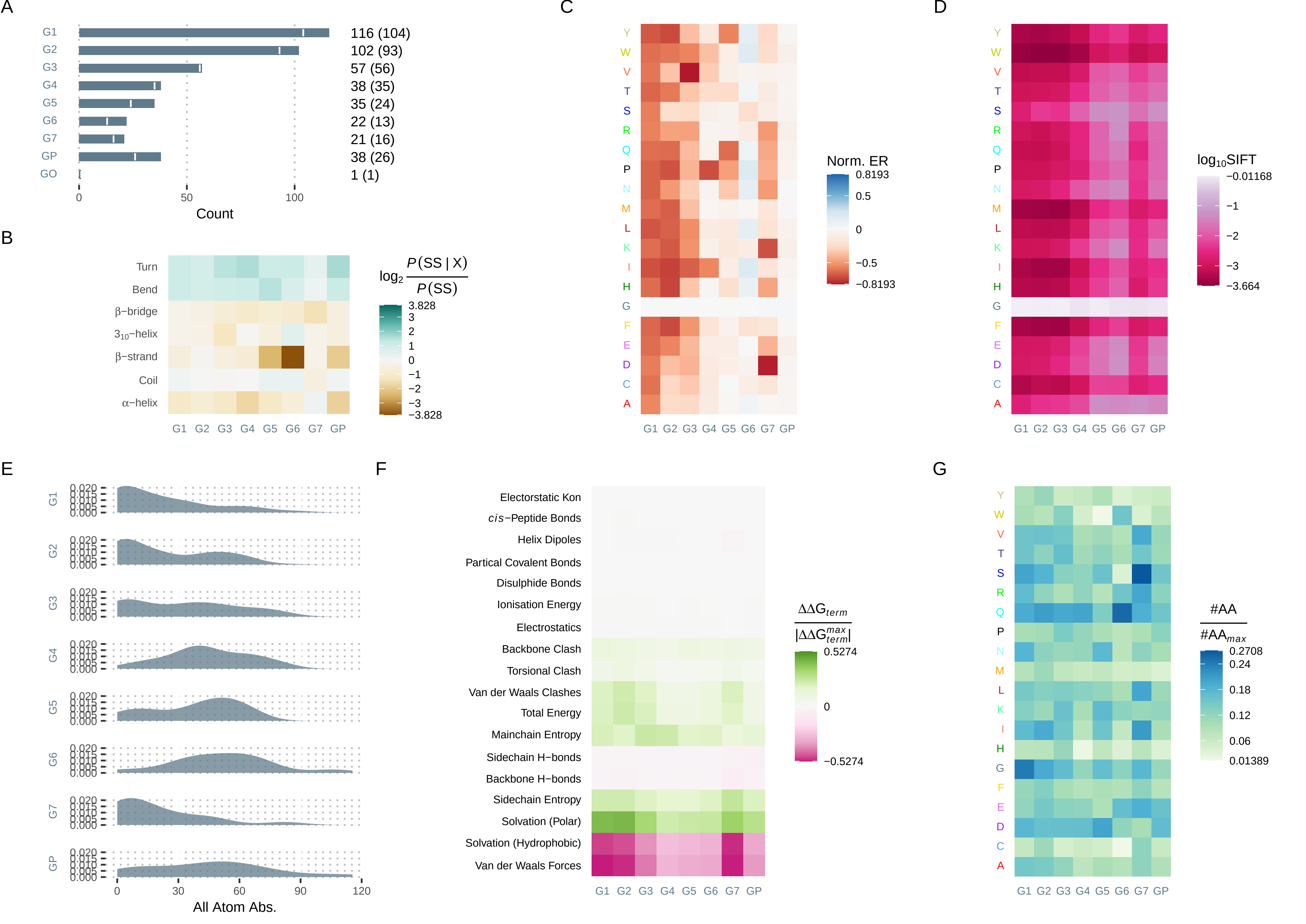

### figureS14.pdf

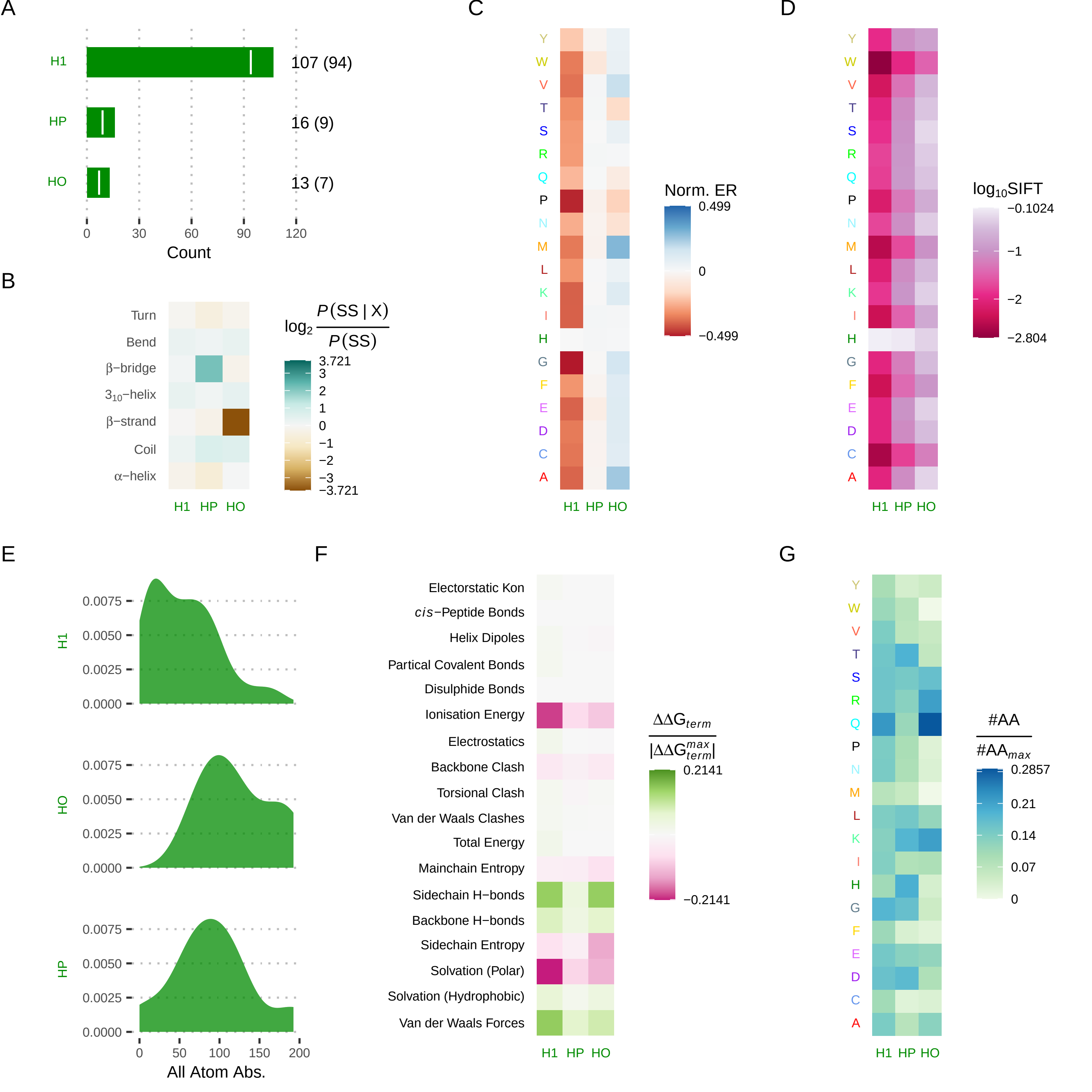

### figureS15.pdf

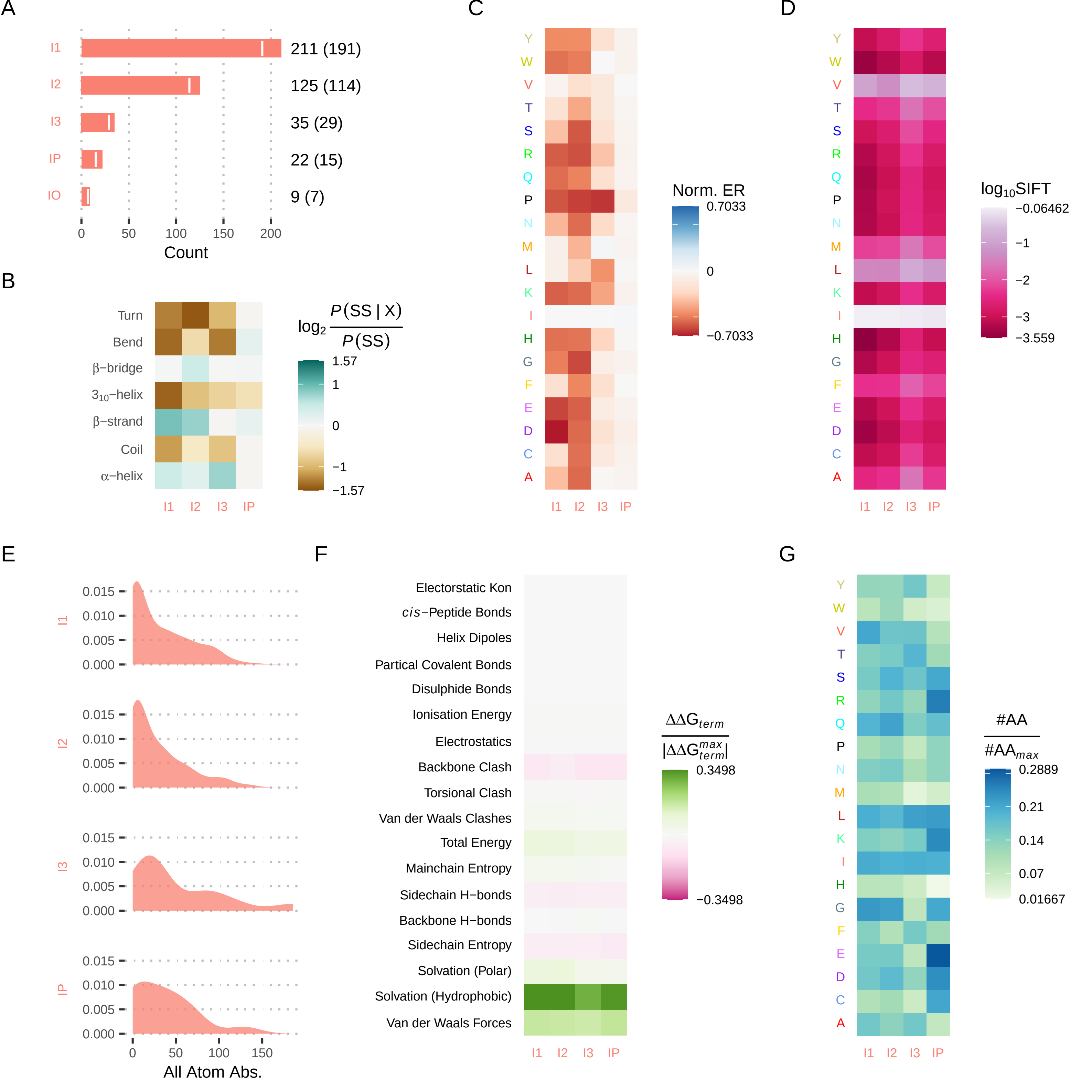

### figureS16.pdf

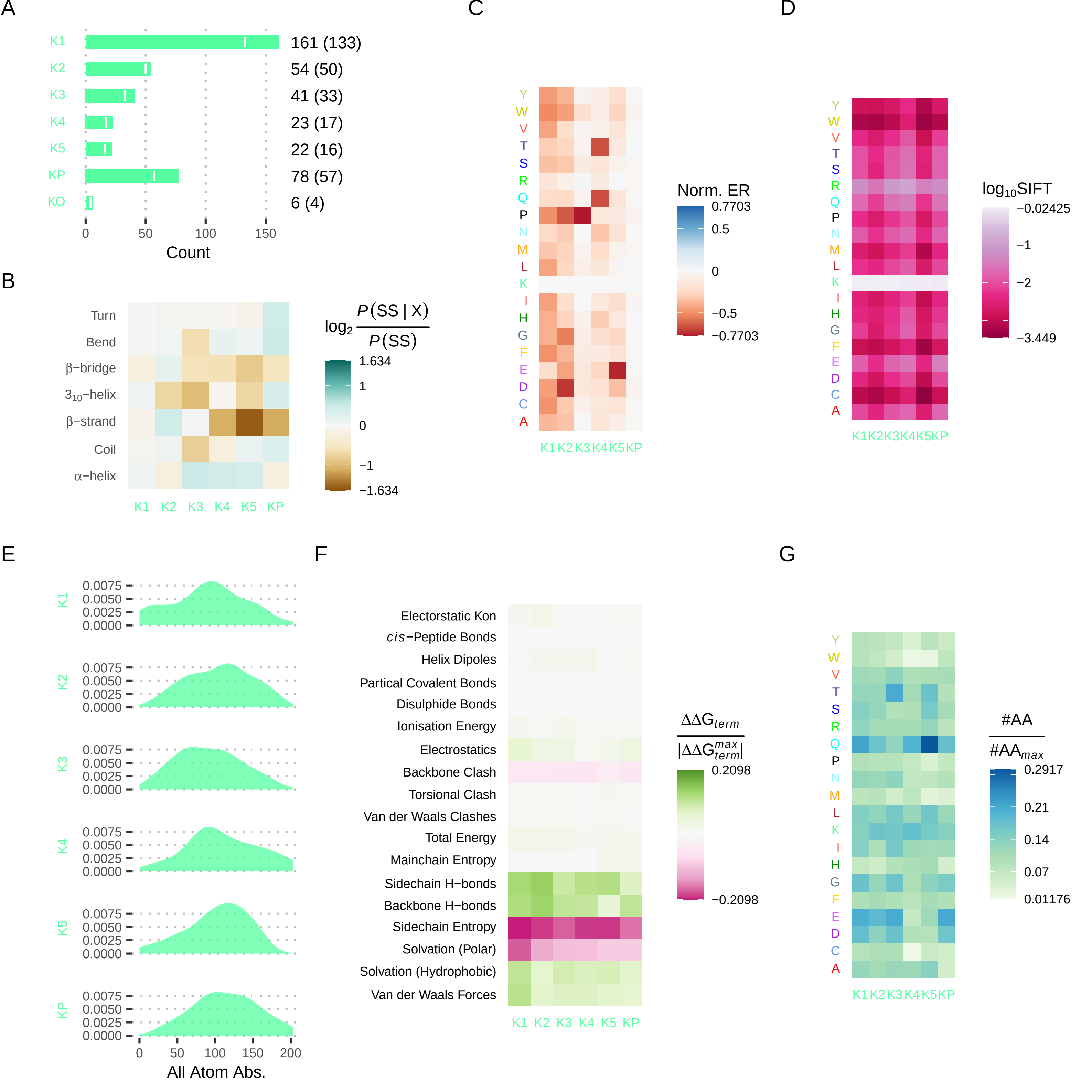

### figureS17.pdf

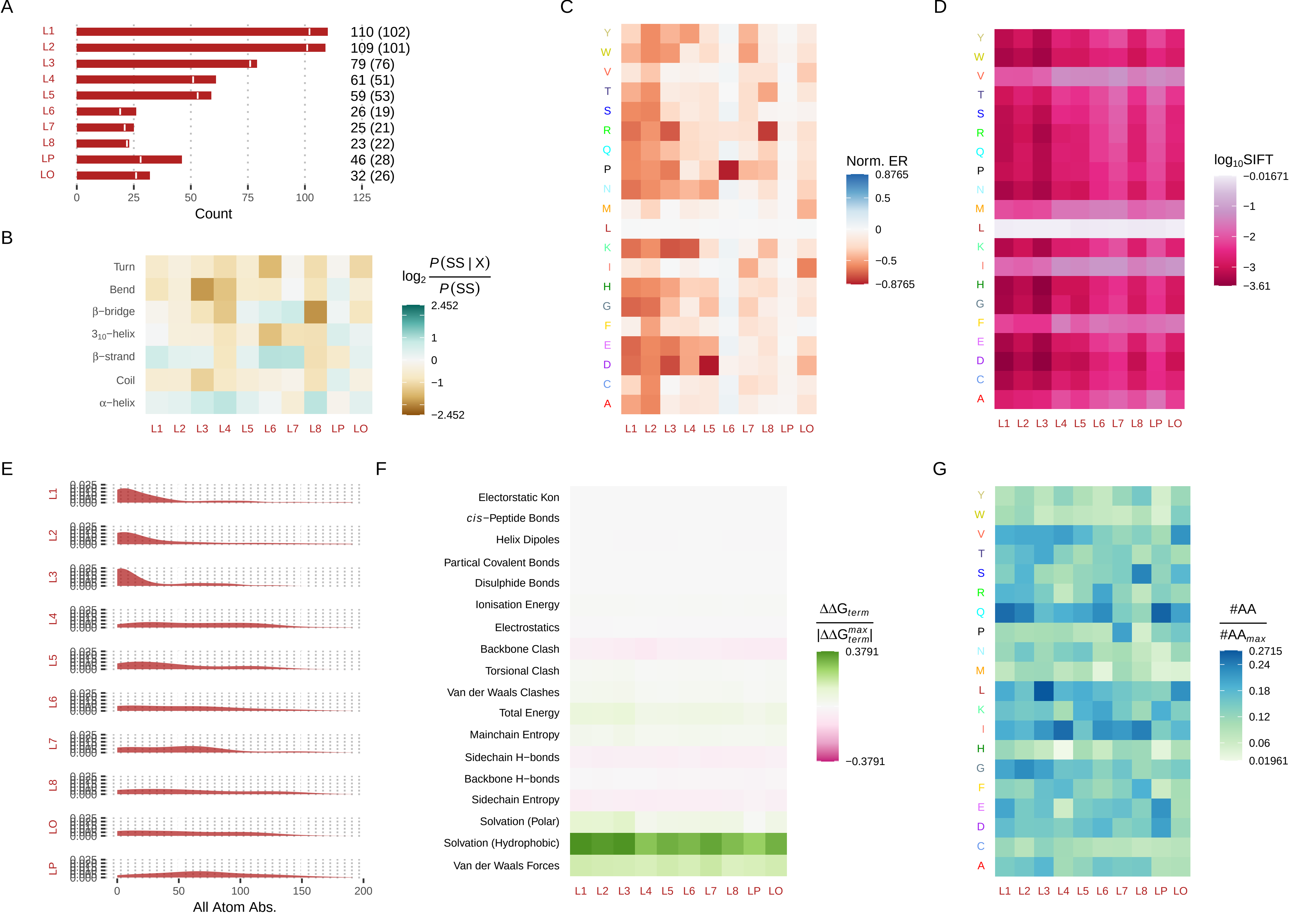

### figureS18.pdf

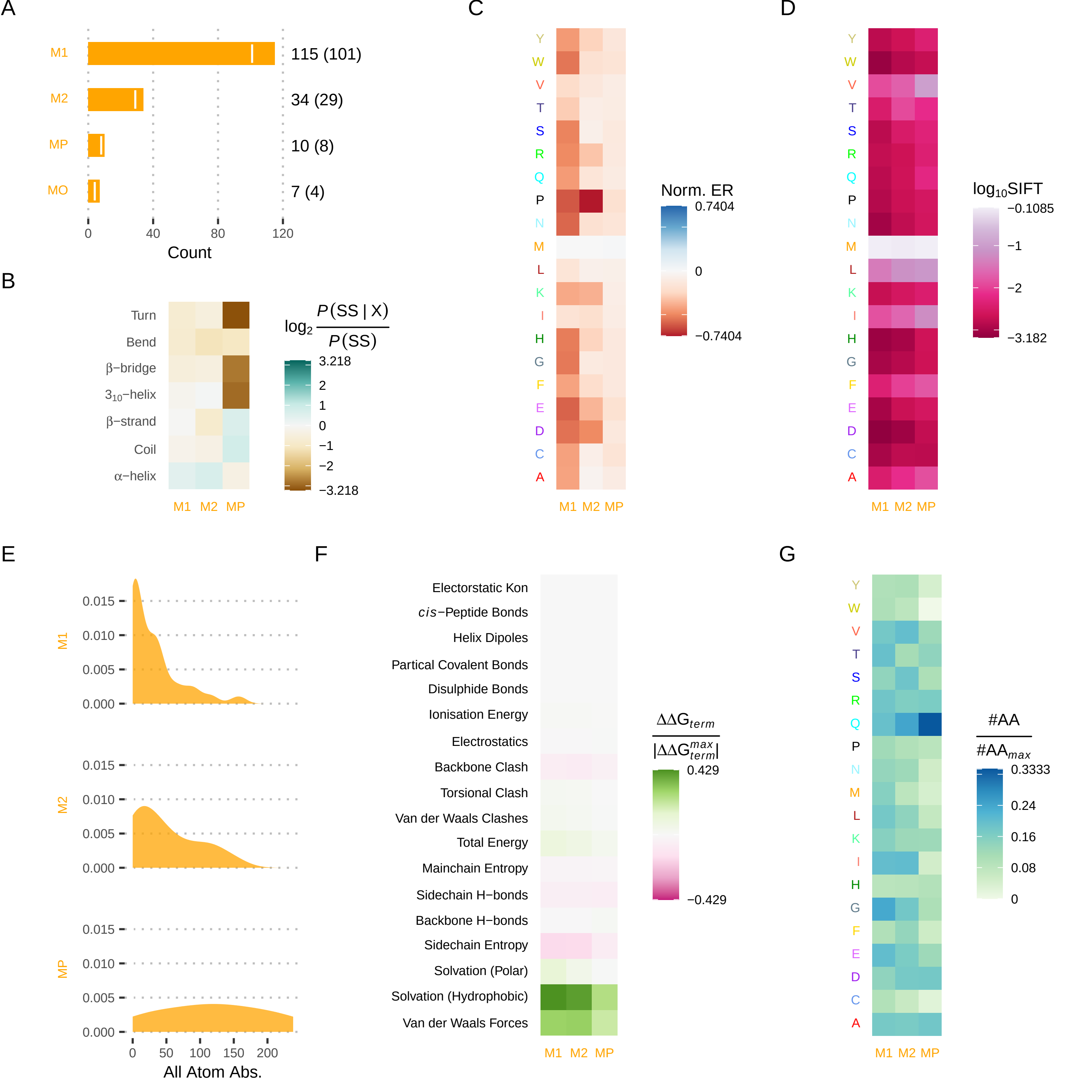

### figureS19.pdf

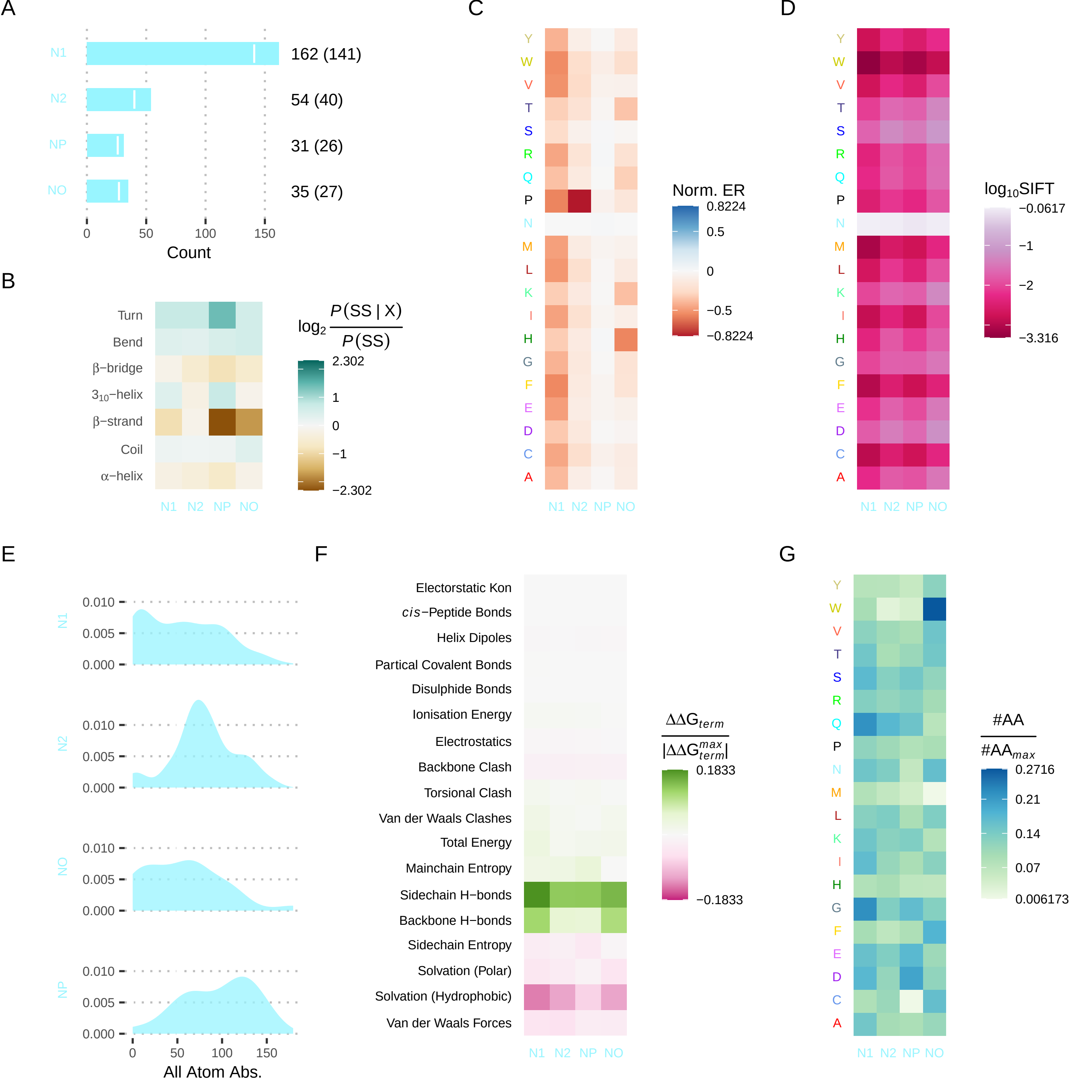

### figureS20.pdf

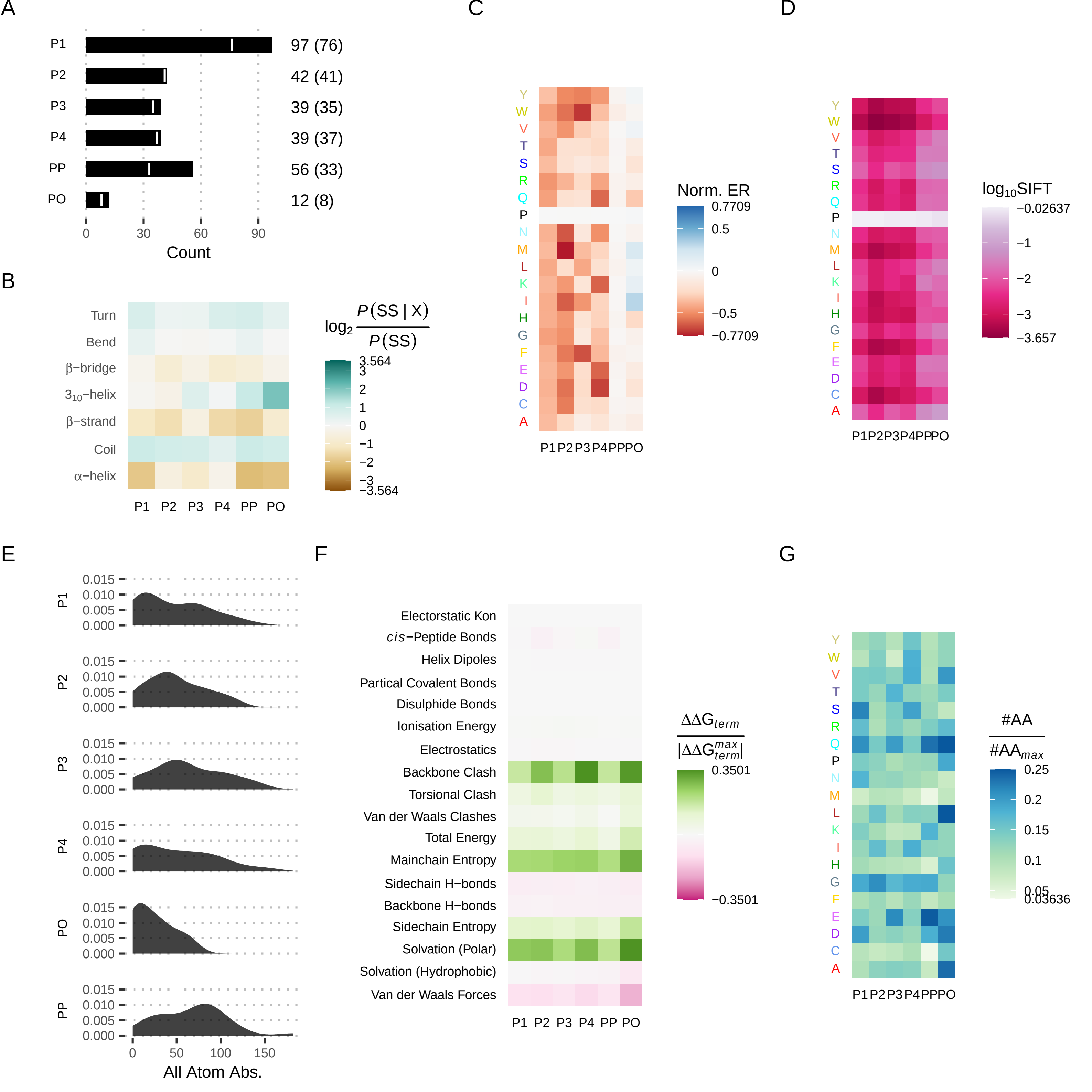

### figureS21.pdf

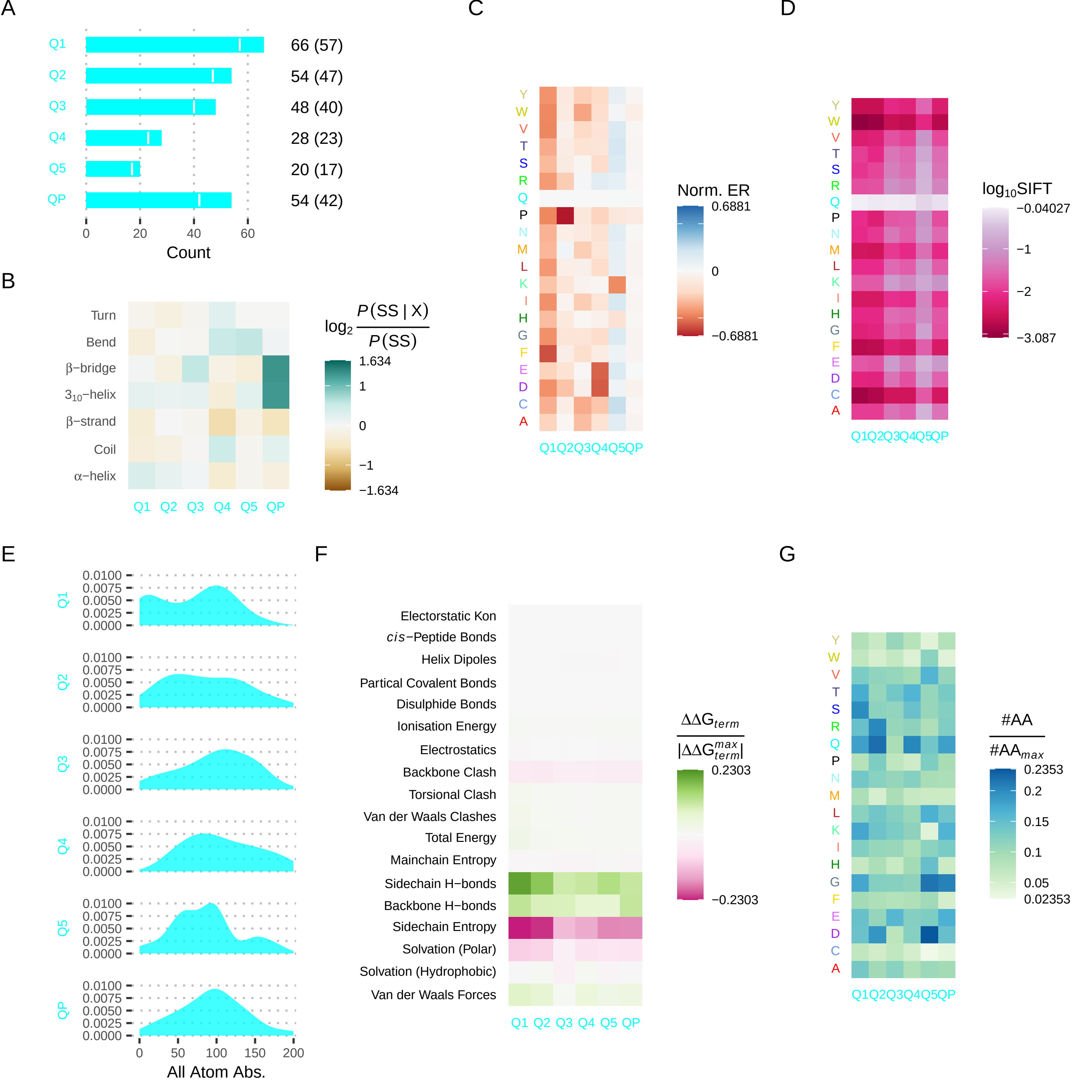

### figureS22.pdf

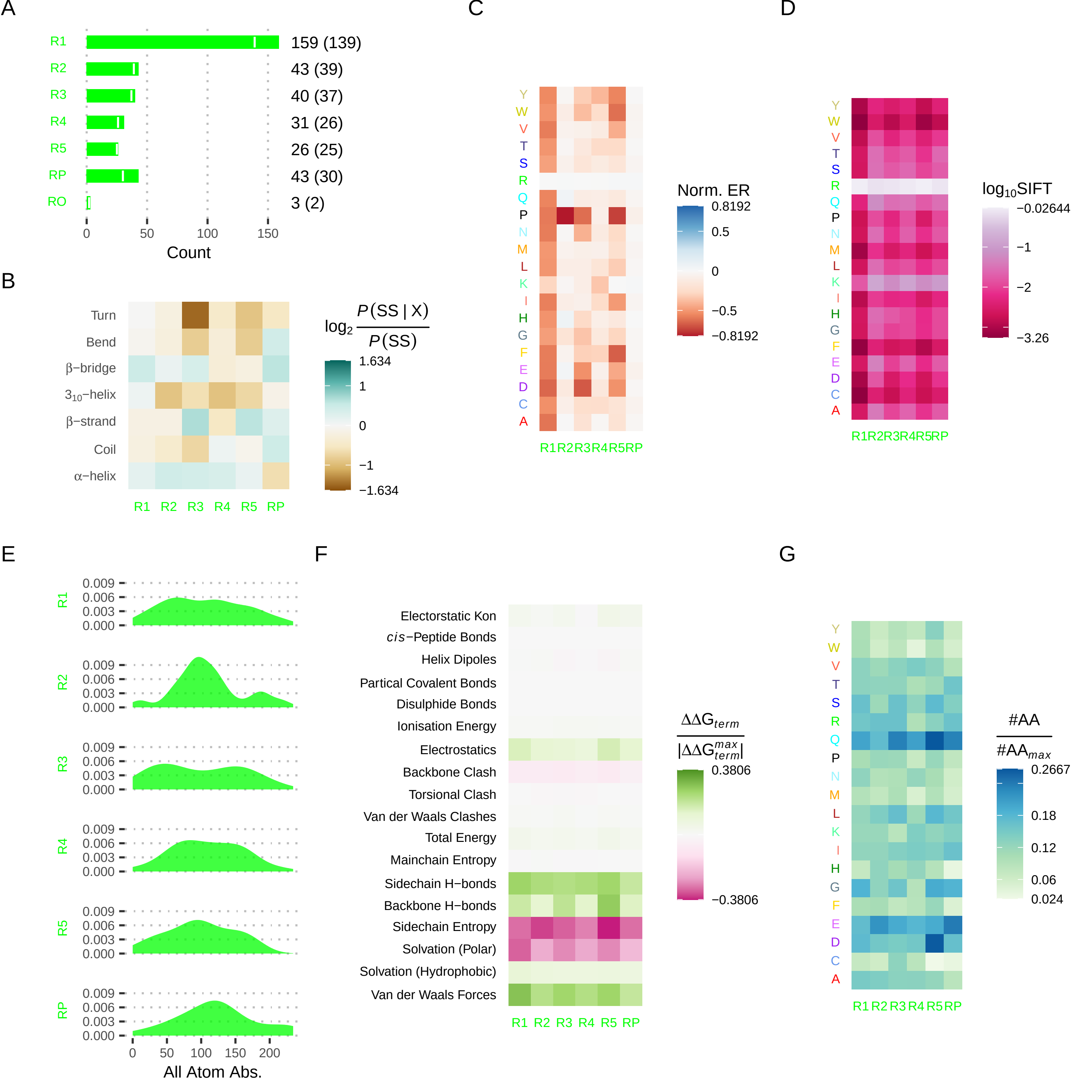

### figureS23.pdf

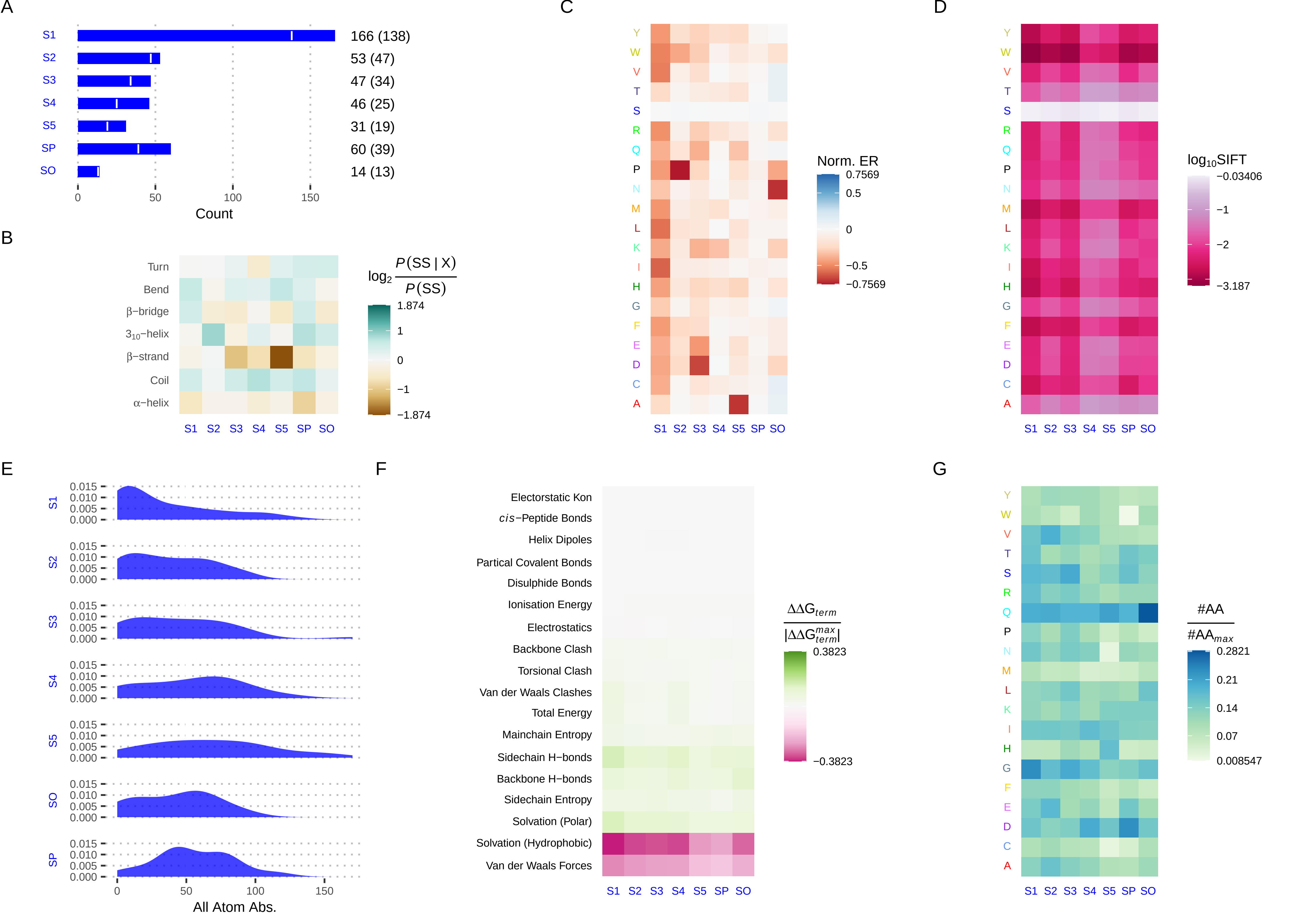

### figureS24.pdf

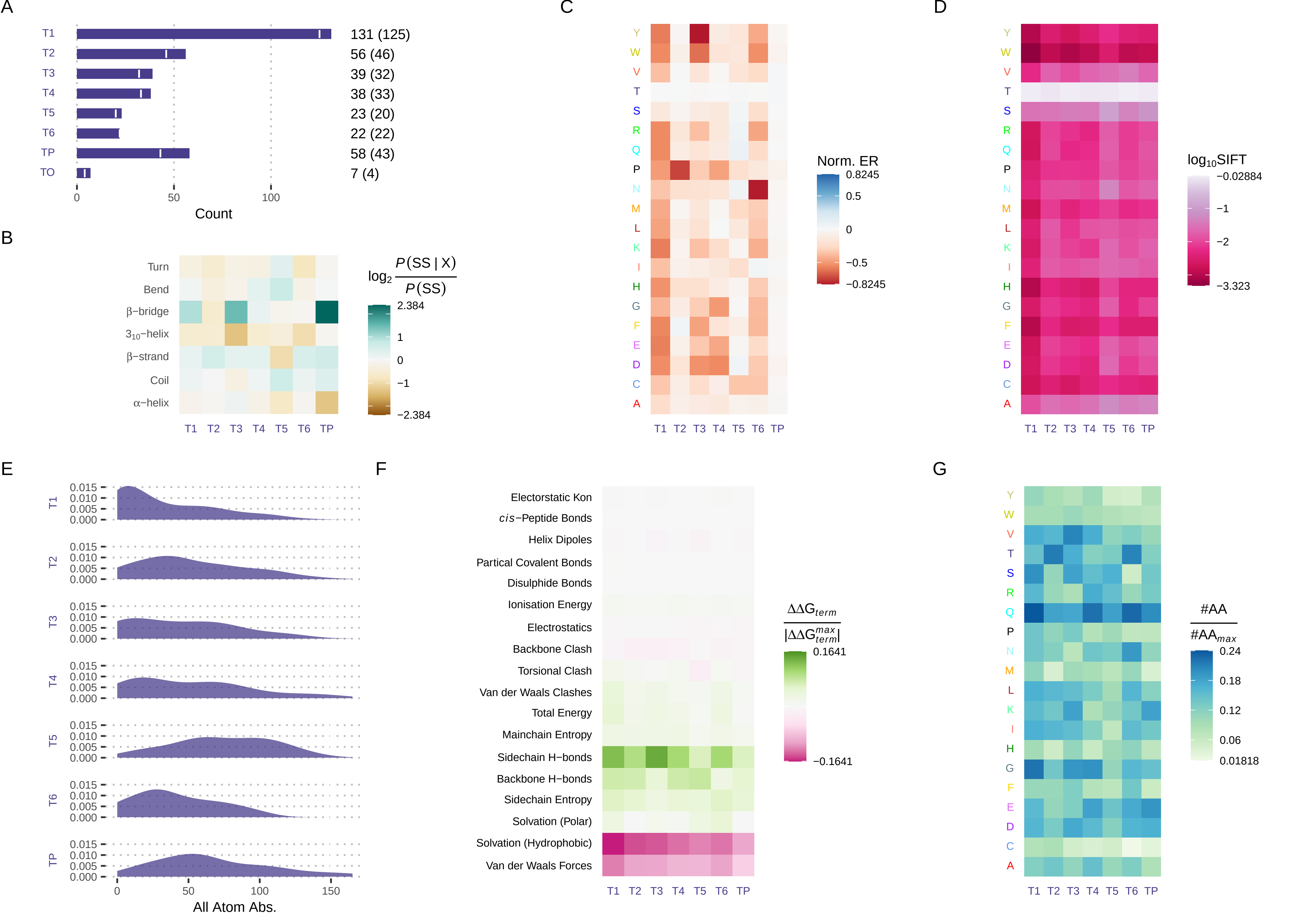

### figureS25.pdf

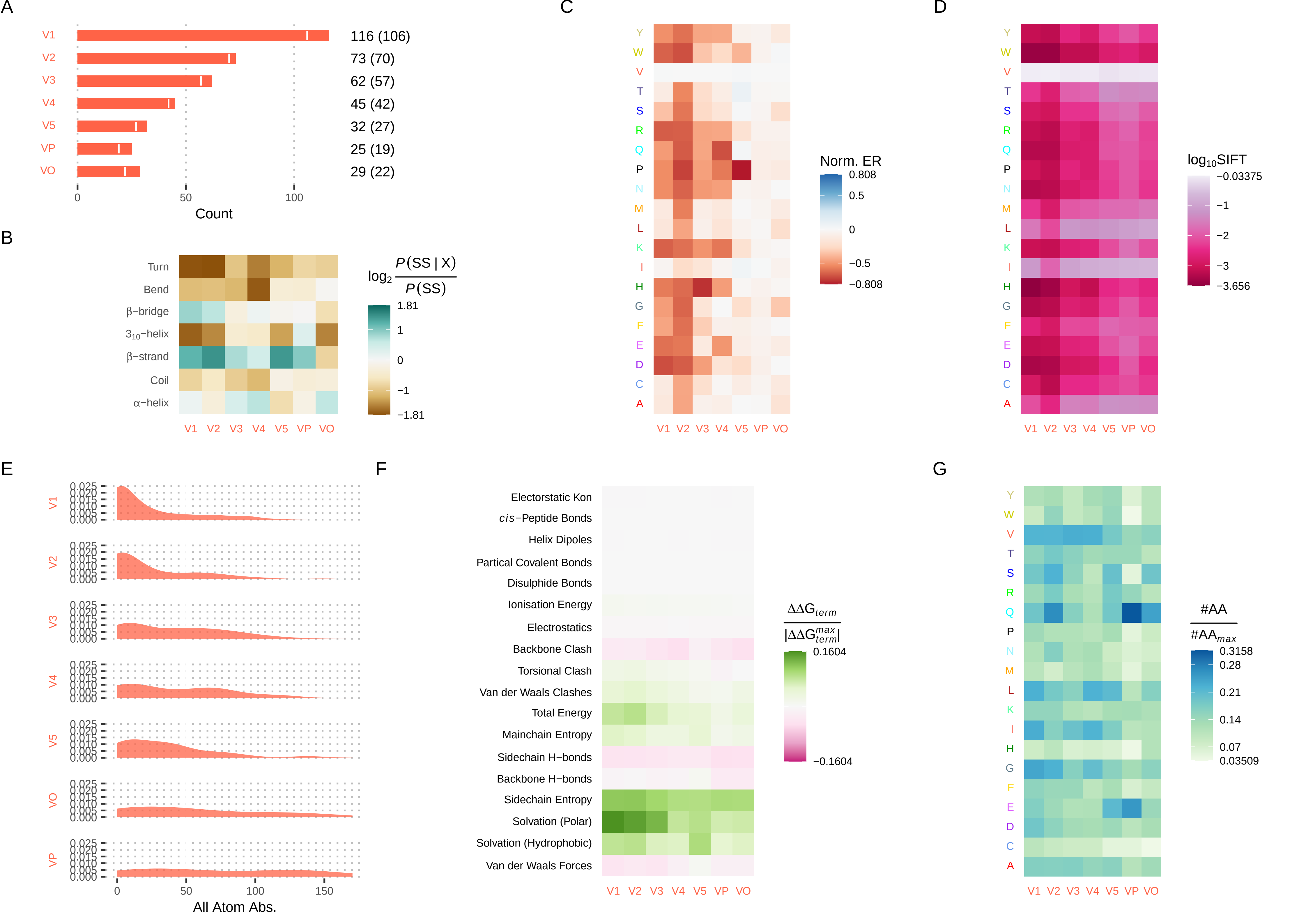

### figureS26.pdf

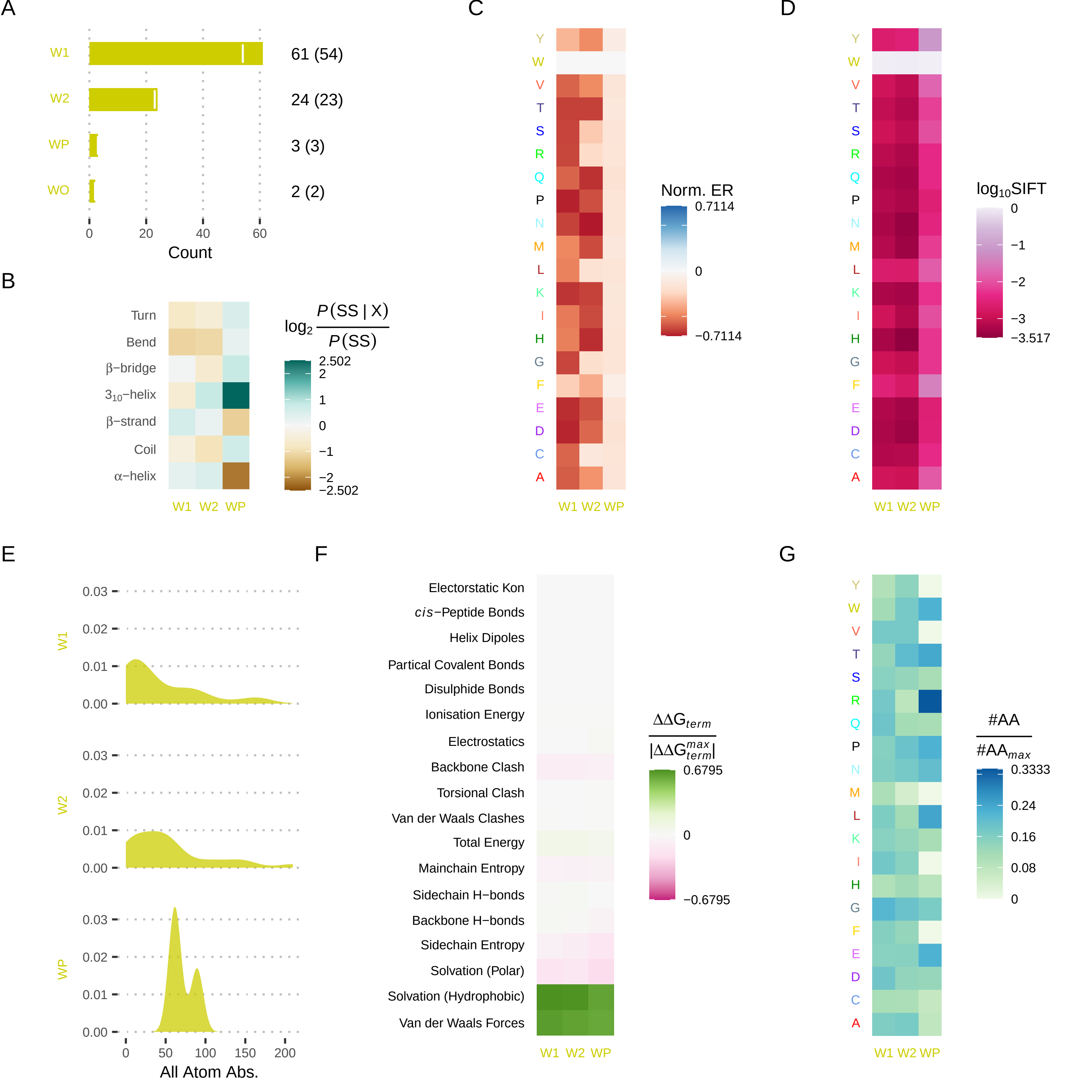

### figureS27.pdf

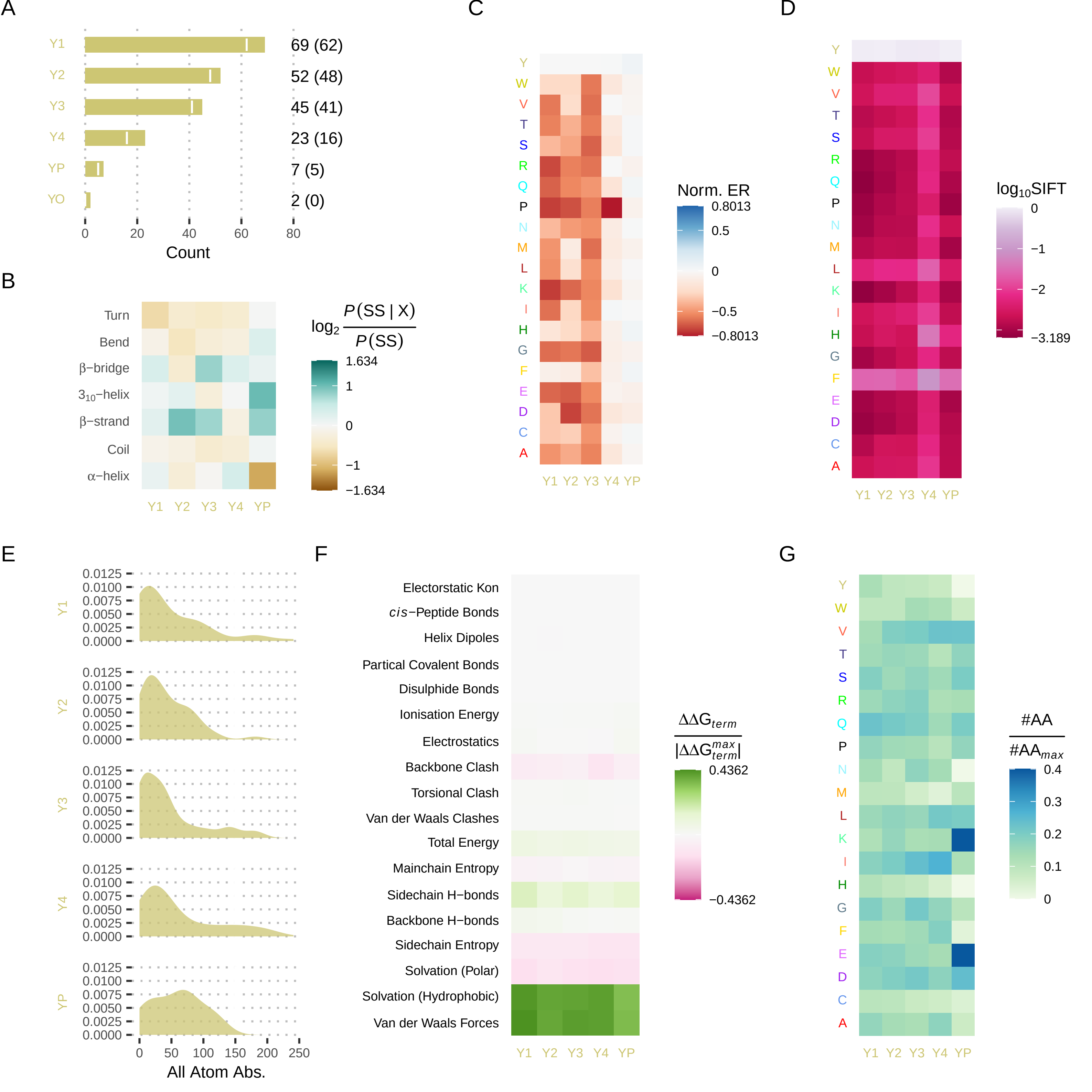
